# Bidirectional communication between nucleotide and substrate binding sites in a type IV multidrug ABC transporter

**DOI:** 10.1101/2025.01.15.633140

**Authors:** Victor Hugo Pérez Carrillo, Margot Di Cesare, Dania Rose-Sperling, Waqas Javed, Hannes Neuweiler, Julien Marcoux, Cédric Orelle, Jean-Michel Jault, Ute A. Hellmich

## Abstract

ATP-binding cassette (ABC) transporters use ATP to transport substrates across cellular membranes. In type IV ABC transporters, including many multidrug resistance (MDR) pumps, communication between the nucleotide-binding domains (NBDs) and transmembrane domains (TMDs) occurs via intracellular coupling helices. However, the precise mechanism of interdomain crosstalk and coordination between the ATP and substrate binding sites remain unclear. Combining nuclear magnetic resonance (NMR) spectroscopy, Hydrogen-Deuterium eXchange Mass Spectrometry (HDX-MS), photo-induced electron-transfer fluorescence correlation spectroscopy (PET-FCS) and functional assays, we identified a conserved cluster of residues at the NBD/TMD interface of the bacterial MDR transporter BmrA. This cluster is crucial for transporter function and relays nucleotide and substrate binding between the two domains via coupling helix 2. Mutations impact both local and global transporter dynamics. Our findings reveal a novel interdomain communication pathway in type IV ABC transporters, shedding light on the intricate coupling mechanisms that enable the coordination of these molecular machines.

## Introduction

ABC transporters are one of the largest protein superfamilies present in all kingdoms of life. They transport chemically diverse substrates including lipids, ions, peptides and vitamins across cellular membranes at the expense of ATP hydrolysis^1,2^. All ABC transporters share a common architecture composed of two structurally diverse transmembrane domains (TMDs) responsible for substrate binding and translocation as well as two cytoplasmic nucleotide-binding domains (NBDs)^2,3^. The four domains can be fused on a single polypeptide (‘full transporter’), or assembled e.g. by two halves each comprising a TMD and an NBD (‘half transporter’) that then dimerize to form a functional transporter. The NBDs are the most sequentially and structurally conserved regions and contain the motifs responsible for ATP binding and hydrolysis, such as the Walker A and Walker B motifs^4–6^. In contrast, the TMDs are structurally and sequentially diverse across the ABC superfamily, resulting in the recent classification into seven subfamilies^3.^

Following this classification, many multidrug resistance (MDR) pumps such as BmrA from *B. subtilis* or mammalian P-glycoprotein (P-gp, ABCB1) belong to the type IV subfamily of ABC transporters^7 8^. In this subfamily, interdomain crosstalk between NBD and TMD is mainly attributed to the proximity of the conserved Q–^9,10^ and X–loop motifs^11–13^ of the NBDs with the coupling helices^14–17^ of the TMDs. In half transporters like BmrA with six transmembrane helices, these intracellular α-helical linkers connect transmembrane helices 2 and 3 as well as 4 and 5. Together with the cytosolic regions of the respective transmembrane helices these linkers constitute the intracellular domains 1 and 2, i.e. ICD1 and ICD2, respectively^18^. Upon assembly into functional transporter dimers, ICD1 interacts with the NBD within the same subunit (in *cis*), while ICD2 reaches over to the NBD of the opposing subunit (in *trans*), inserting into a groove on the NBD surface between the RecA and the α-helical subdomains^12,19^. This ‘swapped topology’ is the hallmark of the type IV ABC subfamily ^3,12^.

Given that ABC transporters efficiently couple ATP hydrolysis with substrate translocation, it is crucial to integrate the existing architectural framework of type IV ABC transporters with a long-range dynamic coupling network that facilitates effective crosstalk between the TMD and NBD. A vast body of work highlighted the importance of the interdomain contacts mediated by the ICDs for the ABC transporter catalytic cycle, including mutagenesis and crosslinking experiments, biophysical and structural as well as computational methods, *e.g.*^12,15–17,20–24^. However, a precise mechanistic understanding of interdomain crosstalk in ABC transporters remains currently amiss, specifically regarding the dynamics of the molecular dialogue between nucleotide and substrate interaction sites that are typically more than 40 Å apart.

Our prior observation that the backbone and sidechain amide chemical shifts of a tryptophan residue remote from the nucleotide binding site in the NBD of the *L. lactis* MDR transporter LmrA responds to the addition of nucleotides^25^ suggested that this residue belongs to a novel long-range sensing network. Therefore, we explored whether homologous transporters such as *E. coli* MsbA or *B. subtilis* BmrA have similar allosteric responses to nucleotides. Here, combining mutagenesis, functional assays, solution ^1^H^15^N and ^19^F NMR spectroscopy, PET-FCS and HDX-MS on the *bona fide* MDR ABC transporter BmrA^8^, which features a conserved type IV subfamily topology^26,27^, we identified a novel communication hinge in the transporter NBD/TMD consisting of three conserved residues in the NBD/TMD interface. In addition to maintaining structural stability, this hinge forms a bidirectional dynamic relay to transmit information on nucleotide and drug interaction between NBD and TMD via ICD2. Our findings thus identify a novel pathway for interdomain crosstalk in a type IV ABC transporter and provide insights into the elaborate interdomain coupling of these sophisticated molecular machines.

## Results

### A conserved cluster of residues in the NBDs of Type IV ABC transporters senses remote nucleotide binding

ABC transporters use ATP to power substrate translocation. The nucleotide interacts with conserved motifs in the NBD, including a lysine residue (K380 in BmrA) from the Walker A motif that assists hydrolysis^28^ (**Fig. 1A**). Nucleotide binding is thought to be relayed to the NBD/TMD interface through another conserved motif, the Q-loop, although the precise molecular pathway remains unknown^9,10^. We previously observed that tryptophan W421 in the isolated NBD of the *L. lactis* MDR ABC transporter LmrA, ∼22 Å away from the nucleotide binding site and upstream of the Q-loop, senses nucleotide binding in NMR chemical shift perturbation experiments^25^. A sequence alignment of type IV ABC transporter NBDs showed that the analogous site typically harbors a bulky hydrophobic side chain such as tryptophan (e.g. BmrA W413, LmrA W421) or leucine (e.g. MsbA L415) whereas in bacterial half transporters closely related to BmrA (i.e. UniProtKB sub-database, 226 sequences extracted from the UniRef50_O06967 database), the tryptophan residue is ubiquitously present (**Fig. 1A, B**).

**Figure 1:**
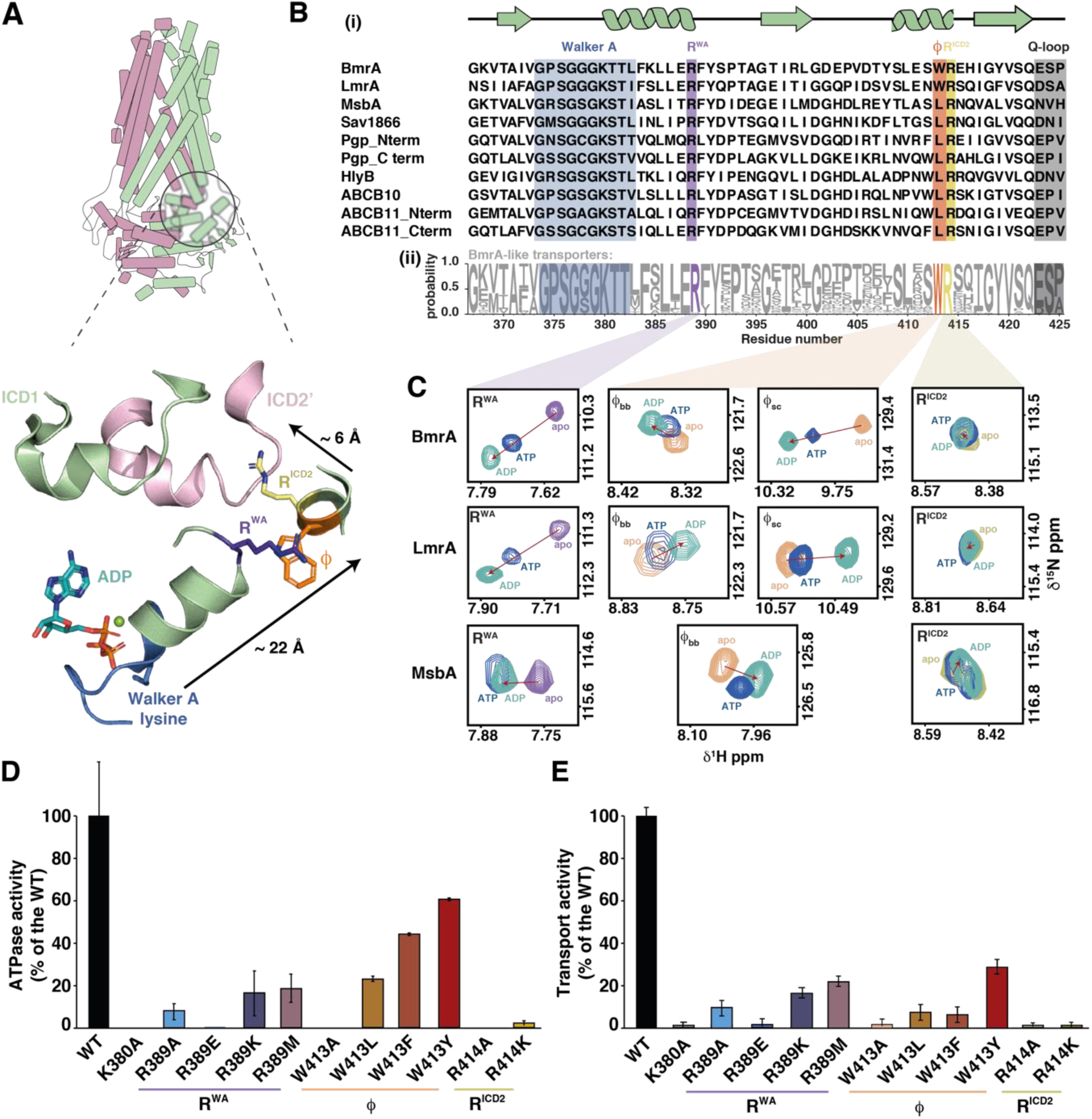
A highly conserved cluster in the NBD of type IV ABC transporter NBDs senses nucleotide binding and is crucial for activity. (**A**) Structure of *B. subtilis* BmrA (PDB ID: 6R81^26^), an archetypical type IV ABC half transporter. The two protomers are shown in pink and green. The zoom shows the Walker A motif (blue) and the Walker A helix leading into the ‘hinge’ consisting of a conserved arginine residue in the Walker A helix (R^WA^, purple), a conserved bulky hydrophobic residue (ϕ, orange) and a conserved arginine pointing towards intracellular domain 2 (R^ICD2^, yellow). See **Supplementary figure 1A** for a structural overlay of this region in other ABC transporters. (**B**) Sequence alignment of the Walker A and hinge region shows a high degree of conservation within the hinge residues (marked purple, orange and yellow) in both human and bacterial type IV ABC transporters (i). The secondary structure is shown on top. Note that in bacterial transporters with at least 50% identity with BmrA, residue ϕ is a strictly conserved Trp residue (ii). For (ii) 226 sequences of type IV ABC transporter NBDs from the UniRef50_O06967 database were used and logo sequence was created using WebLogo3^31^. Residue number is based on the sequence of *B. subtilis* BmrA. (**C**) Nucleotides are sensed by the hinge region. Comparison of NMR spectra of ^15^N-labeled NBDs of BmrA, LmrA and MsbA in the apo state (purple, orange or yellow resonances, respectively) and in the presence of 10 mM ATP (blue) or ADP (teal) reveals chemical shift perturbations of the backbone (and in the case of tryptophan also side chain, ϕ_sc_) amide resonances of the three hinge residues. Shown is a zoom into the respective ^1^H, ^15^N-HSQC NMR spectra (**Supplementary figures S1B, C**) to highlight the three hinge residues. (**D, E**) The hinge residues are crucial for transporter function. ATPase activity of BmrA variants reconstituted in MSP1E3D1 nanodiscs prepared with *E. coli* total lipid extract (D) and fluorescence-based transport assay with doxorubicin in inside-out vesicles prepared from *E. coli* cells overexpressing BmrA variants (E). In both cases, values were normalized to the WT set as 100%. Results shown are the mean of three biological triplicates with three technical replicates each. Protein expression and folding were not affected by the mutations (**Supplementary Fig. S3**)

To elucidate a possible function for this residue, we compared the fingerprint ^1^H, ^15^N-HSQC solution NMR spectra of the 29 kDa ^15^N-labeled NBDs of three homologous bacterial type IV transporters: the multidrug transporters *B. subtilis* BmrA and *L. lactis* LmrA, as well as the *E. coli* lipid A transporter MsbA in the apo and the nucleotide bound states. To this end we took advantage of our previously published and *de novo* determined NMR backbone assignments for the three proteins (BMRB entries 17660 ^29^, 51156 ^30^ and 52626) (**Fig 1C)**.

In all three NBDs, the chemical shift of the abovementioned hydrophobic residue (BmrA W413, LmrA W421, MsbA L415) responded to the addition of ATP or ADP (**Fig 1C, Supplementary figure S1**). In the case of the Trp residues in BmrA and LmrA, a shift in both the backbone and sidechain amide resonances was observed. To reflect the chemical nature of the side chain, we denote this residue as ϕ. It resides in a short α-helix within the RecA subdomain and is consistently followed by an arginine (BmrA R414, LmrA R422, MsbA R416), which points towards the coupling helix from ICD2 of the *trans* protomer in these half transporters (**Fig. 1A)**. Facing residue ϕ, the C-terminal tip of the helix that harbors the Walker A (WA) motif and its conserved lysine residue contains another highly conserved arginine residue (BmrA R389, LmrA R397, MsbA R391). Both arginine residues, hereafter referred to as R^ICD2^ and R^WA^ to mark their respective location, also respond to the addition of ATP or ADP, an effect particularly pronounced for residue R^WA^ (**Fig. 1B, C, supplementary Fig. 1**).

The three conserved residues appear to be ideally positioned to serve as a ‘communication hinge’ between the nucleotide binding site and the TMD (**Fig. 1A**). Together, R^WA^, ϕ and R^ICD2^ may form a functional nexus in the NBD, allosterically linking the nucleotide binding site to the NBD/TMD interface. This connection likely extends from the Walker A motif and the following helix through R^WA^, to ϕ, and into R^ICD2^ reaching the TMD.

### The hinge is important for protein stability and transporter function

Located centrally in the NBD, the hinge plays a role in both nucleotide sensing and structural integrity. CD spectroscopy and analytical SEC revealed that point mutations in the hinge of BmrA, LmrA and MsbA destabilize the isolated NBDs, with the most pronounced effects for mutations of residue R^WA^ (**Supplementary figure S2)**. Notably, substitutions of BmrA W413 with leucine and alanine could not be purified in the context of the isolated NBD. Furthermore, the melting temperatures (*T_m_*) of the hinge mutant NBDs that could be obtained were consistently lower than the respective wildtype (WT) proteins (**Supplementary Table 1)**, although nucleotide dissociation constants were only slightly altered for most cases (e.g. *K_D_, _ADP_* = 214±43, 221±105, 221±54 and 266±97 µM for BmrA NBD WT, R398K, W413F and R414K, respectively). In contrast, more drastic substitutions such as R398M and R414A reduced the affinity of two- and ten-fold, respectively (**Supplementary Table 2).**

To explore the role of the hinge for transporter function, point mutations were introduced into full-length *B. subtilis* BmrA to yield the respective BmrA R^WA^ (R398A/K/E/M), ϕ (W413A/F/Y/L) and R^ICD2^ (R414A/K) variants. As a negative control for functional assays, the Walker A lysine mutant BmrA K380A^32^ was also prepared. The successful expression and purification of all hinge variants in detergent micelles suggests that the TMD can partially mitigate folding defects that were pathological for the isolated NBD (**Supplementary Fig 3A-C**). However, similar to the isolated NBDs, a consistent decrease in the *T_m_* for the hinge variants was observed in the full-length transporter (**Supplementary Table 3)**. Notably, while the WT and the other hinge mutants were stabilized by nucleotides, both R414 substitutions and the K380A mutant failed to do so, suggesting that they are not able to reach the outward facing (OF) state (**Supplementary Table 3**).

The ATPase activity of purified BmrA full-length WT, K380A and hinge mutant constructs was determined in detergent micelles, lipid nanodiscs and liposomes (**Fig. 1D, Supplementary Fig 3D, E**). Transport activity was measured in inside-out vesicles prepared from cells overexpressing BmrA variants using the fluorescent substrate doxorubicin (**Fig. 1E; Supplementary Fig 3F, G**). Mutations in the hinge severely impaired both ATPase activity and transport, even for conservative substitutions. Mutation of the R^ICD2^ residue to alanine or lysine completely abolished function. R^WA^ and ϕ residue mutants showed a more graded response: The R389A/E and W413A mutants showed very little to no activity, while the R389K/M and W413L mutants displayed some and the W413F/Y mutants retained substantial residual ATPase activity. Intriguingly, the W413F and W413Y mutants appeared functionally uncoupled, with ATPase activity much less affected than doxorubicin transport. Consistent with our findings, W413 mutations in BmrA were also reported to differentially impair Hoechst 33342 transport (W413A>W413F>W413Y), although the ATPase activities of the mutants were not measured^33^.

### Nucleotide binding site and communication hinge are bidirectionally coupled

ABC transporters coordinate nucleotide binding and hydrolysis in the NBD with substrate binding and translocation in the TMD. We hypothesized that the hinge may act as a bidirectional signal relay between nucleotide and substrate binding sites.

To test this, we first examined how a mutation in one of the three hinge residues affected the remaining two, taking advantage of the NBD variants that were sufficiently thermostable for NMR spectroscopy (**Supplementary Table S1**). 2D ^1^H, ^15^N-correlation and 1D ^19^F-NMR spectra showed that that single hinge residue mutations caused chemical shift perturbations and in some cases line broadening for the resonances of the remaining two hinge residues (**Fig. 2A, B**, **Supplementary Fig. 4**). This suggests both structural and dynamic local coupling within the hinge.

**Fig. 2:**
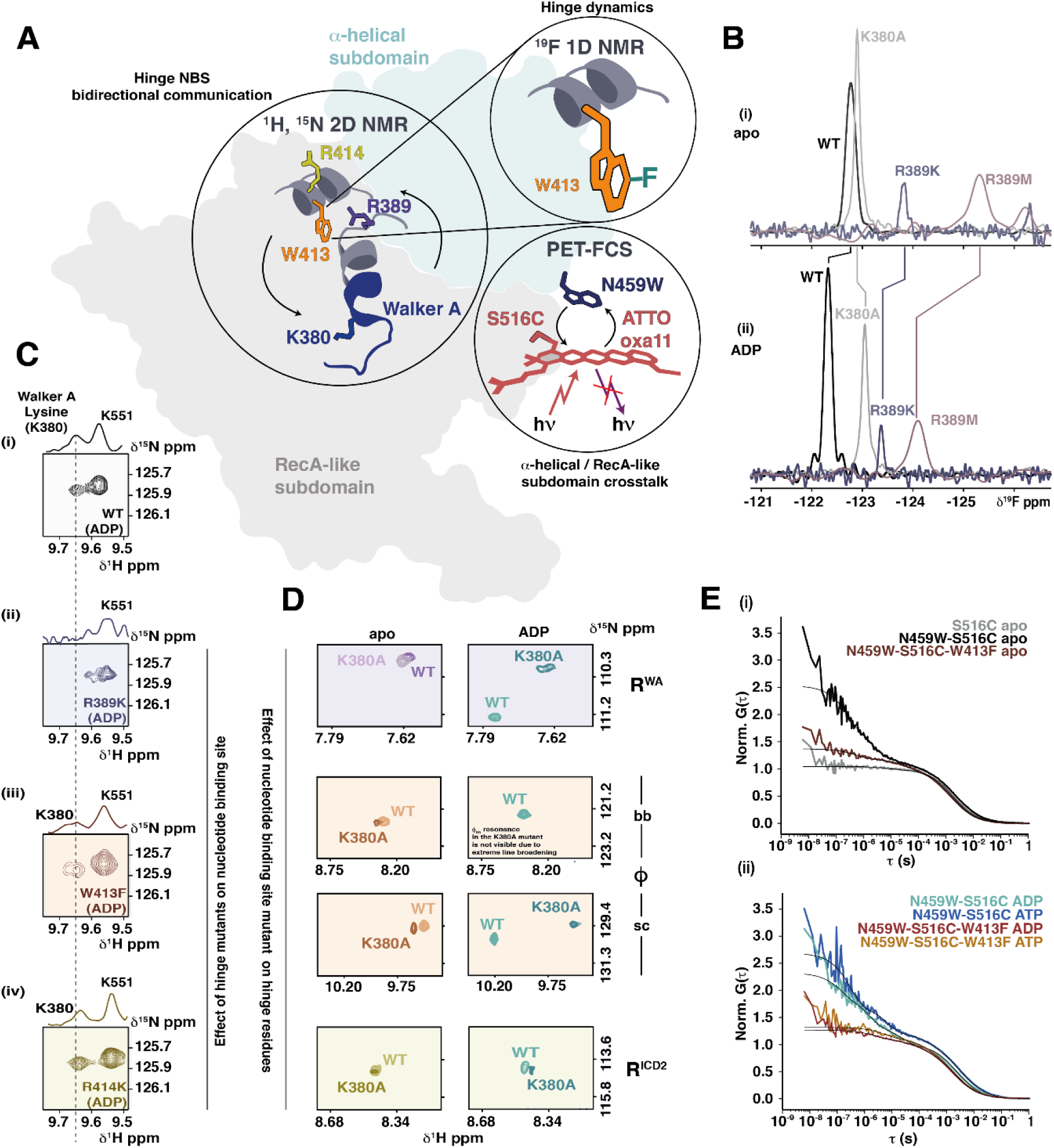
Bidirectional coupling between hinge and nucleotide binding site in the BmrA NBD. (**A**) Complementary methods (^19^F 1D NMR, ^1^H, ^15^N 2D NMR and PET-FCS) were used to explore the bidirectional crosstalk between the nucleotide binding site and hinge within the BmrA NBD, as well as intradomain structural rearrangements caused by hinge mutations. (**B**) 1D ^19^F NMR spectra of ^19^F-5Trp-labeled BmrA NBD variants in the absence (i) or presence (ii) of nucleotides. Residue ϕ, W413, is the only native tryptophan in the BmrA NBD, thus giving rise to a single fluorine resonance in the ^19^F NMR spectrum of WT BmrA NBD (black). Mutation of the R^WA^ residue into lysine (blue) or methionine (mauve) or the Walker A lysine K380 to alanine (grey) led to chemical shift changes and line broadening of ϕ, suggesting an influence on the ϕ sidechain dynamics. Mutations of residue R^ICD2^ led to fully aggregated protein and were thus not included. Spectra of BmrA NBD WT (250 µM), K380A (300 µM), R389K (100 µM) and R389M (250 µM) constructs were recorded at 298 K at 600 MHz with 512, 256, 4096 and 10240 scans, respectively. (**C**) Hinge mutations are sensed by the conserved Walker A lysine, K380. Shown is a zoom into the ^1^H, ^15^N-HSQC and the corresponding 1D projection for the K380 resonance in the WT (i), R398K (ii), W413F (iii) and R414K (iv) constructs. In the apo state, the resonance of K380 shows severe line broadening, thus experiments were carried out with 10 mM ADP. (**D**) Mutation of the Walker A lysine is sensed by the hinge residues. Depicted are zooms into the ^1^H, ^15^N-HSQC spectra of the WT and K380A BmrA NBD in the absence (left) and presence of 10 mM ADP (right) monitoring the backbone NH resonances of residues R^WA^ (top) and R^ICD2^ (bottom). For residue ϕ, both the backbone (bb) and sidechain (sc) NH resonances are shown (middle). (**E**) Mutation of the hinge alters intradomain NBD dynamics. PET-FCS autocorrelation functions of fluorescently labeled BmrA NBD constructs were recorded in the absence (i) or presence of nucleotides (ii). PET-FCS reporters were designed to monitor fluctuations between α-helical and RecA domain. In a cysteine-less background (BmrA NBD C436S), the fluorescently modified S516C, S516C-N459W and S516C-N459W-W413F served as a control, a reporter for hinge motions, and a probe for the role of conserved W413, respectively. Black lines are fits to the data using a model for a single diffusion containing one, two or three single-exponential relaxations.

For instance, mutation of residue R^WA^ to methionine or lysine resulted in the complete loss of signal for the indole side chain NH resonance of BmrA W413 in the ^1^H, ^15^N HSQC spectra (**Fig. 2B**). To nonetheless obtain information about the residue ϕ sidechain in the context of the R^WA^ mutant, residue ϕ (W413) was fluorinated to introduce an alternative NMR reporter. This can be easily achieved by feeding 5F-tryptophan to the expression host^34–36^. Conveniently, residue ϕ is the only native tryptophan residue in the BmrA NBD, yielding a single resonance in the 1D ^19^F NMR spectrum (**Fig. 2B (i)**). Of note, ^19^F NMR measurements could only be carried out for the WT and R^WA^ mutants, since R^ICD2^ mutants became unstable with the fluorine label and could not be purified. Mutation of neighboring residue R^WA^ led to line broadening and a strong chemical shift for the tryptophane side chain of ϕ, reflecting altered sidechain dynamics. Importantly, the fluorinated protein retained its sensitivity to nucleotides. Introduction of the Walker A lysine mutant K380A mutation also induced a chemical shift change in the ^19^F resonance of residue ϕ, both with and without nucleotide (**Fig. 2B (ii)**), strongly supporting the notion of direct communication between the hinge and the nucleotide binding site.

Conversely, we then investigated how hinge mutations affected the nucleotide binding site using the backbone NH resonance of K380 as a reporter. Since K380 is inherently dynamic, its ^1^H, ^15^N backbone amide is only visible in the ADP-bound state^30^ (**Fig. 2C(i)**). Mutation of any of the three hinge residues led to chemical shift changes or severe line broadening for the K380 backbone amide resonance, with the most pronounced effects from R^WA^ mutations, followed by ϕ and R^ICD2^ substitutions (**Fig. 2C (ii)-(iv), Supplementary Figure S5**). Finally, mutating the Walker A residue to alanine impacted the backbone (and in the case of ϕ also the sidechain) NH chemical shifts and line widths of all three hinge residues, with more pronounced effects in the presence of 10 mM nucleotide than in the apo state, despite the reduced affinity of the K380A mutant for ADP (**Fig. 2D**, **Supplementary Fig. S6, Supplementary Table 2)**. These findings suggest nucleotide dependent, bidirectional crosstalk between the nucleotide binding site and hinge in the ABC transporter NBD.

### Hinge mutation affects intra-NBD dynamics between α-helical and RecA-like domain

An ABC transporter NBD consists of an α-helical and a RecA-like subdomain, whose dynamics are modulated by nucleotide binding^37^. We investigated whether a hinge mutation influences these intradomain dynamics using photoinduced electron transfer fluorescence correlation spectroscopy (PET-FCS), which measures structural fluctuations of biomacromolecules on the ms to ns timescale detected through fluorophore quenching by a nearby tryptophan sidechain^38,39^.

To label the BmrA NBD for PET-FCS, we first generated a construct lacking the native cysteine residue, BmrA NBD C436S, which facilitated site-directed thiol modification with a fluorophore and does not affect BmrA function^27^. A newly introduced cysteine residue, S516C in the RecA domain, was then covalently labeled with an ATTO Oxa11 maleimide fluorophore. Combined with an engineered tryptophan residue in the α-helical domain, N459W, this allowed to monitor conformational fluctuations across the subdomain cleft (**Fig. 2A, E, Supplementary Table S4**). As a control, the corresponding autocorrelation function (ACF) of a construct lacking the tryptophan in the α-helical domain was well described by diffusion of a globular protein through the confocal detection focus that was void of additional fluorescence fluctuations in the sub-ms time regime (**Figure 2E(i), gray trace**). The observed, additional 8 µs relaxation had a negligible amplitude of only 5% (**Supplementary Table S4**). This confirmed that detected changes by FCS indeed originated from quenching interactions between the engineered Trp at position N459W and the Atto Oxa11 label. In contrast, the ACF of the fluorescently labeled protein carrying the S516C-N459W mutation pair exhibited three single-exponential sub-ms relaxations of substantial amplitude originating from intramolecular subdomain dynamics (**Figure 2E(i), black trace**), i.e. corresponding to three modes of motions of the two domains relative to each other on the µs-ns time scale. Nucleotide binding further modulated the subdomain kinetics (**Figure 2E(ii)**). In the construct with an intact hinge, ADP accelerated the 80-µs mode of motion to 24 ±4 µs, while binding of ATP slowed the motion to 253 ±16 µs (**Figure 2E(ii), Supplementary Table S4**).

Mutation of the hinge residue ϕ (W413F) altered the intradomain NBD dynamics substantially (**Figure 2E(i), brown trace**). One mode of motion, i.e., the 2.5 µs relaxation, was abolished and the amplitude of the ns relaxation was considerably reduced, while the 80 µs relaxation was accelerated. In the mutant, nucleotide binding slowed the 39-µs relaxation to time constants >100 µs and the 0.57 µs motion to 2.7 ± 0.2 and 5.7 ±0.7 µs with ATP and ADP, respectively (**Figure 2E(ii), Supplementary Table S4**).

While it is difficult to assign specific relaxations to specific motions, which may include shearing or breathing, the PET-FCS results unambiguously show that the interaction of either nucleotides or the mutation of the hinge alter intradomain dynamics of the NBD through a pathway of allosteric communication.

### The communication hinge impacts global transporter dynamics through ICD2

Hydrogen–deuterium exchange coupled to mass spectrometry (HDX–MS) and ^19^F NMR spectroscopy are powerful tools for studying membrane protein conformational dynamics^24,34,40,41^. Here, we combined these methods to investigate whether the hinge affects interdomain communication between the NBD and TMD by comparing full-length BmrA WT and the W413F mutant (**Fig. 3, Supplementary Fig. S7 S8 S9**). The W413F hinge mutant was chosen because it retained substantial ATPase activity (∼50%) while the transport activity was severely impaired (<10%) (**Fig. 1, Supplementary Fig. S4D, E).** This allowed to trap the transporter with MgADP_*_V_i_ to emulate the transition state for ATP hydrolysis.

**Fig. 3:**
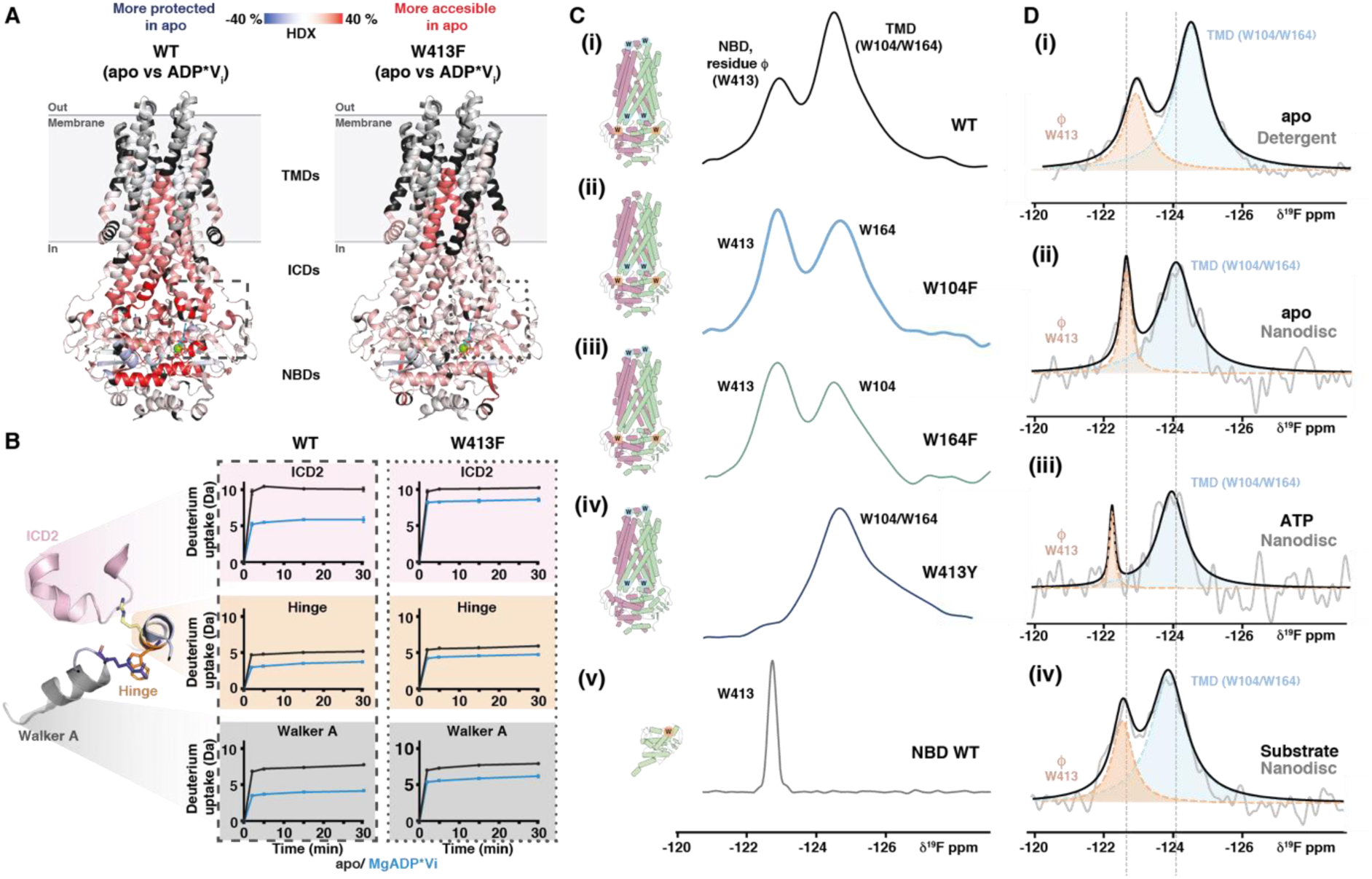
The hinge impacts global transporter dynamics and senses both nucleotide and substrate binding. **(A)** Hydrogen–deuterium exchange coupled to mass spectrometry (HDX–MS) on BmrA WT and BmrA W413F in nanodiscs comparing deuteration in outward and inward facing (OF/IF) states. Shown are the differences in HDX after 30 min between the apo and the MgADP*Vi trapped state (after incubation of 10 mM MgATP and 1 mM V_i_) mapped on the cryoEM structure of BmrA in the OF state (PDB: 7OW8, nucleotide shown as cyan sticks, Mg^2+^ as green sphere). Peptides with no significant deuteration differences (p-value = 0.05, see Methods) are shown in light gray and uncovered peptides in black. Peptides in red represent regions that are more accessible in the apo state than in the ADP*Vi state (See **Supplementary Fig. 7** for sequence coverage and absolute deuteration plots). (**B**) The differential deuterium uptake between the apo and MgADP*Vi states is subdued in the vicinity of the hinge upon introduction of the W413F mutation. Shown is a zoom into the hinge region highlighting differences in HDX between WT (left) and hinge mutant (right). Selected significant peptides (p value<0.05) from ICD2 (amino acid residues 216 – 236; VKASNAEDVEYGRGKMGISSL), the hinge (amino acid residues 413 – 427; WREHIGYVSQESPLM) and the Walker A helix (amino acid residues 363 – 383; VIEAGKVTAIVGPSGGGKTTL) all display reduced HDX in the MgADP*Vi trapped state (blue) compared to the apo state (black). These differences are more pronounced for the WT than the hinge mutant. (**C**) 1D ^19^F NMR on detergent solubilized ^19^F-5Trp-labeled BmrA. BmrA contains three native tryptophan residues (W104, W164, W413) that were individually mutated to obtain the respective ^19^F resonance assignments ((i)-(iv)). The spectrum of the isolated NBD carrying only native tryptophan W413 confirms the correct assignment (v). (**D**) Nucleotide and substrate binding affect the hinge. The ^19^F NMR assignments in the detergent state (i) could be directly transferred to ^19^F-5Trp labeled BmrA reconstituted in MSP1E3D1 nanodiscs with *E. coli* polar lipids (ii). Addition of ATP (iii) or the substrate reserpine (iv) resulted in changes in chemical shift and linewidths for the ^19^F resonance of W413. For better visibility, we carried out a spectral deconvolution using a pure Lorentzian–line shape fitting.

Overall, > 90% peptide sequence coverage of BmrA WT and W413F reconstituted in nanodiscs either in the apo state or trapped states could be achieved in the HDX experiments (**Supplementary Fig. S7A**). In line with previous studies^42,43^, the NBDs had better peptide coverage than the TMDs, which are partially shielded from proteolysis by the MSP-lipid belt.

In the apo state, the WT protein exhibited significant flexibility, particularly within the NBDs and ICDs (**Supplementary Fig. S7C)**. Upon MgADP_*_V_i_ trapping, HDX globally decreased in the WT transporter showing rigidification, a behavior consistently seen across type IV ABC transporters upon nucleotide binding ^8,11, 44–47^ (**Fig. 3A**). Deuterium exchange differences between apo and trapped states were most pronounced for the conserved motifs in the NBD (e.g. Walker A and C-loop) and the intracellular domains 1 and 2 (ICD1, ICD2), particularly the C-terminal end of TMH4, leading into ICD2, and TMH5, which transitions from ICD2 into the TMD (**Fig. 3A**, red regions, **Fig. 3B**).

In comparison to the WT, the W413F hinge mutant exhibited subdued exchange differences between apo and MgADP_*_V_i_ trapped states in key regions, including the Walker A motif, the hinge and ICD2 and the surrounding helices TMH4 and 5 (**Fig. 3A** right, **Fig. 3B, Supplementary Fig. S7**). Notably, a major consequence of the W413F mutation was increased HDX in the ADP/Vi trapped state compared to the WT (**Fig. 3B** and **Supplementary Fig. S7B-D**), highlighting that nucleotide-dependent interdomain communication between NBD and TMD is primarily mediated via ICD2, and is significantly disrupted when the hinge is mutated.

### Lipid environment and substrates influence hinge dynamics

To complement the HDX-MS data on the hinge region, which provides information at peptide level, we used ^19^F NMR spectroscopy in solution with the full-length transporter. In addition to the hinge residue W413, BmrA contains two other native tryptophan residues: W104 in ICD1 and W164 located within the extracellular loop between TMH3 and TMH4 (**Fig. 3C, Supplementary Fig. 8A**). Using three single point mutants targeting each of the native tryptophan residues in the full-length transporter and the isolated NBD with only W413, we unambiguously assigned the three fluorine resonances of ^19^F-5Trp labeled BmrA to the two tryptophan residues in the TMD and to W413 in the NBD (**Fig. 3C**). Structural and functional integrity of the fluorinated transporter was confirmed by CD spectroscopy, transport and ATPase assays (**Supplementary Fig. 8B-D**).

Importantly, the fluorine resonances were readily transferable from detergent-solubilized to nanodisc-reconstituted protein (**Fig. 3D (i), (ii)**). While the W413 resonance became narrower, indicating altered NBD dynamics in the lipid environment as reported for other type IV ABC transporters^48,49^, it remained distinct from the TMD tryptophan residues, suggesting that both the global NBD dynamics, as well as local hinge dynamics are impacted by the transporter environment.

Next, we investigated the effect of nucleotide or substrate binding on W413. As seen in the ^19^F NMR spectra of the isolated NBD (**Fig. 2B**), ATP addition led to chemical shift changes and line narrowing (**Fig. 3D (iii)**). Substrate addition led to severe line broadening of the W413 ^19^F NMR signal, showing that substrates directly affect hinge dynamics (**Fig. 3D (iv)**). Due to the changes in pH associated with adding a saturating amount of doxorubicin to the NMR samples, which led to sample precipitation, we instead used reserpine, another BmrA substrate^8,11^. To nonetheless obtain information on the transporter dynamics with doxorubicin, we carried out HDX-MS in the presence of doxorubicin, achieving similar peptide coverage as for the apo and ATP_*_V_i_ trapped transporter (**Supplementary Figure 9**). Comparison of BmrA WT in the absence and presence of doxorubicin showed decreased exchange in the TMD, notably TMH4, 5 and ICD2 in the presence of the drug (**Supplementary Figure 9B**). This shows that substrate binding decreases dynamics in regions also seen to become rigidified by nucleotide addition (**Fig. 3A**). A similar trend was seen for the W413F hinge mutant, although generally substrate-evoked effects were noticeably subdued compared to the WT, suggesting reduced coupling as also apparent from the activity assays (**Supplementary Figure 9**, **Fig. 1D, E)**.

In summary, HDX-MS and ^19^F NMR demonstrate that nucleotide and drug binding to NBD and TMD respectively influence hinge dynamics, and that hinge mutations globally affect protein dynamics and interdomain crosstalk.

## Discussion

ABC transporters are multidomain membrane proteins that rely on precise coordination between NBDs and TMDs to drive ATP-dependent substrate transport. We thus hypothesized that structural ‘nodes’, i.e. amino acid residues or regions that respond to both nucleotide and substrate binding, enable bidirectional and interdomain crosstalk. Using sequence alignments, functional assays, solution NMR, HDX-MS and PET-FCS on the bacterial MDR transporter BmrA, an archetypical type IV ABC transporter, we identified three highly conserved residues that influence transporter stability, ATPase and transport activity, thus acting as a functional nexus in the NBD/TMD interface.

Structurally, these residues belong to a hinge connecting the nucleotide binding site with the drug binding site. Ligand binding at either end of this L-shaped pathway affects the dynamics of the three residues (**Fig. 4A**). Likewise, mutation of the hinge residues impacts ATP and substrate interactions, demonstrating bidirectionality of this crosstalk. The Walker A-motif is at one extremity of this pathway. Mutation of the invariant lysine (K380 in BmrA) was shown to prevent the transition from the inward to the outward facing state^32,42^. Interestingly, the consequence of the mutation of R414 is redolent of that observed with the K380 mutants supporting that the conformational changes is propagated through the L-shape connecting pathway proposed here.

**Figure 4:**
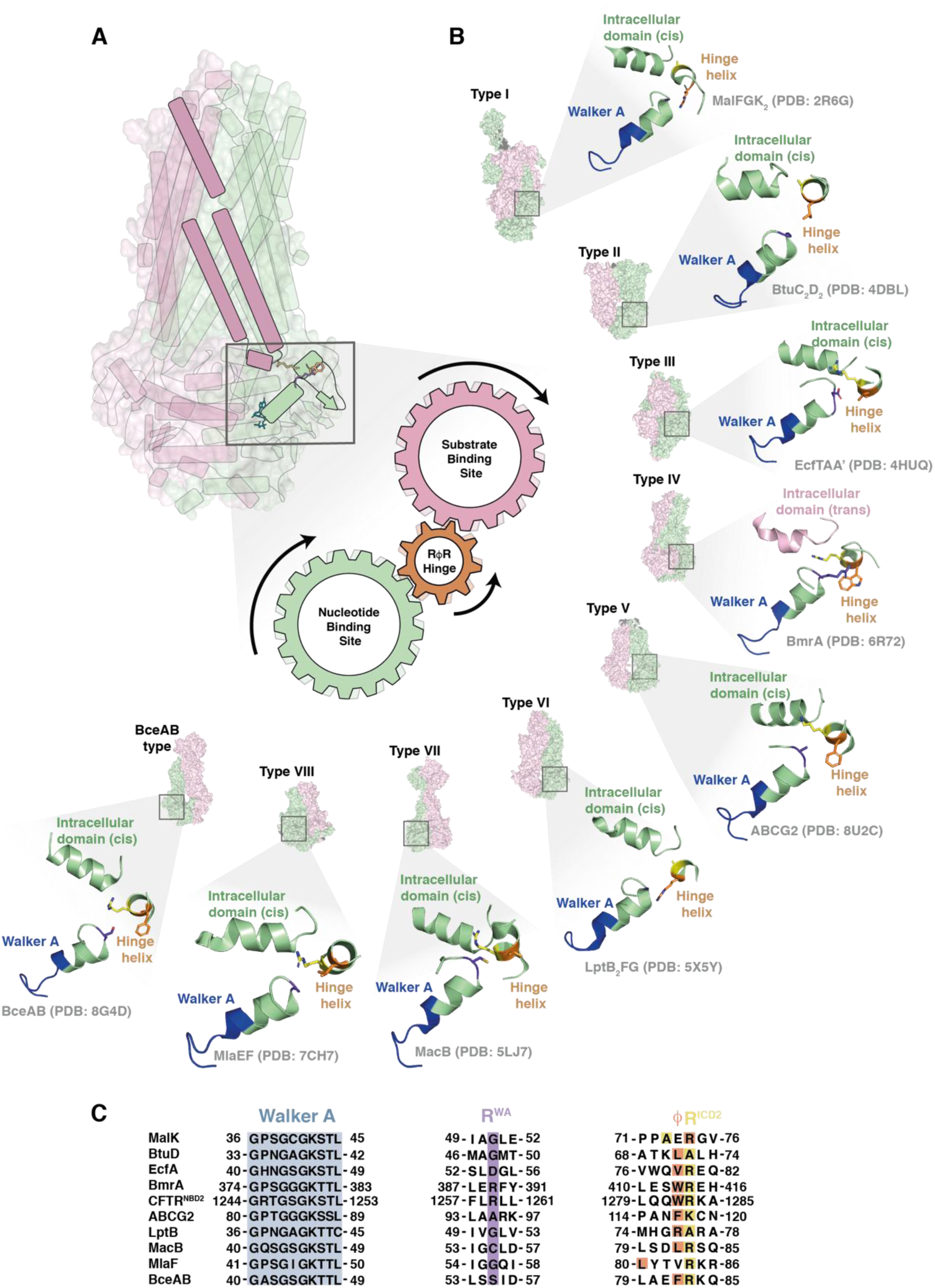
Common features of the hinge region across ABC transporter families. **(A)** The hinge connects nucleotide and substrate binding site. Like cogwheels, the residues enable crosstalk via ICD2. (**B**) Comparison of the core elements of the hinge in different familes of ABC transporters. Residues R^WA^-ϕ−R^ICD2^ are shown in orange, Walker A motif in blue and the intracellular domain in green (cis) or pink (trans). ABC transporter families I-VIII and BceA are presented by surface depictions of representative members and a zoom into the respective hinge region, i.e. MalFGK_2_ (PDB: 2R6G ^62^), BtuC_2_D_2_ (PDB: 4DBL ^63^), EcfTAA’ (PDB: 4HUQ ^64^), BmrA (PDB: 6R81 ^26^), ABCG2 (PDB: 8U2C ^65^), LptB_2_FG (PDB: 5X5Y ^66^), MacB (PDB: 5LJ7 ^67^), MlaEF (PDB;7CH7 ^68^) and BceAB (PDB: 8G4D ^69^). (**C**) Sequence alignment of the Walker A and hinge region for the ABC transporters shown in (B) was carried out using Clustal Omega^70^. Note that in some cases, the sequence alignment was adapted to reflect the 3D structure of the respective proteins (see main text for details). In LptB_2_FG and MalFGK_2_, an inversion of the properties of the sidechains in the positions ϕ and R^ICD^ is seen and that in MalFGK_2_ and MlaEF residues ϕ and R^ICD^ are not contiguous.

Large scale conformational changes and structural rigidification upon ligand binding are a hallmark of allosteric coupling in type IV transporters^11,24,50^. Indeed, BmrA becomes less dynamic upon substrate or ATP binding as seen with HDX-MS. It is conceivable that the hinge acts as “dynamic buffer”, thereby enabling the cushioning of and directed transmission of ligand-induced conformational changes throughout the protein. Possibly, this enables the protein to interact with structurally diverse substrates, which may impose different structural demands on the pathways connecting NBD and TMD. This buffering function is compromised in hinge mutants, disrupting the delicate balance required for coupling. Since mutations can uncouple ATPase and transport function, it is indeed long-range information flow, rather that the inherent functionality of the protein, that seems to be compromised when the hinge is disturbed.

In addition to ligand binding, the hinge may also contribute to the intriguing lipid dependence of ABC transporter function^51^. Adding ABC transporters to the growing list of membrane proteins for which ^19^F solution NMR spectroscopy has been successfully applied^34,52–54^, we used it on detergent solubilized and nanodisc-reconstituted BmrA. Nucleotides, drugs and the membrane environment all affected the dynamics of hinge residue W413, suggesting that in addition to bidirectional intramolecular crosstalk, the hinge may also integrate exogenous cues such as membrane composition or fluidity.

Overall, substitutions of residue ϕ were generally better tolerated than mutations of either of the two conserved arginine residues (**Fig. 1**). The unique ability of arginine to mediate numerous different interactions, including hydrogen bonds, salt bridges, cation-π and π-π interactions between its guanidinium group and aromatics, as well as hydrophobic interactions may play a decisive role here^55,56^. While conservative substitutions of R^WA^ maintained at least some residual activity, even the conservative substitution to lysine for R^ICD2^ was not tolerated and led to an inactive transporter. In structures of BmrA, the sidechain of R^ICD2^ points towards ICD2 **(Supplementary Figure 10**). Together, the structural and functional data suggest that the three hinge residues may act like interlocked cogwheels, connecting the NBS to hinge, and the hinge to the substrate interaction sites and vice versa (**Fig. 4A)**

Importantly, the hinge seems to be pivotal to stabilize the transition state of the transporter and plays a role both in the intra-NBD (between RecA- and α-helical domain) as well as in the global dynamic response of the protein to nucleotide and substrate binding. While both drug and ATP binding decreased WT protein flexibility, this was less pronounced with hinge mutations, which in some cases were apparently not able to reach the outward open state (i.e. R414 mutants, see above). In line with studies on MsbA^14,16^, another type IV ABC transporter, our results show that while both ICD1 and ICD2 are crucial for driving the conformational changes of the catalytic cycle, hinge mutations primarily affect ICD2. Within the family of type IV transporters, the importance of the hinge identified here in BmrA may thus be of general relevance, despite some sequence variability (**Fig. 4B**). For instance, in CFTR (cystic fibrosis conductance regulator, ABCC7), NBD2 features a QWR sequence, while in NBD1, a glycine residue is found in position R^WA^, and the WR residues in ϕ and R^ICD2^ are present but not contiguous. Nonetheless, mutations causing cystic fibrosis are found in the hinge, including a deletion mutant adjacent to the residue homologous to BmrA R398 and at positions corresponding to W413 and R414^57^. Moreover, the side chain of residue F508 points towards the CFTR hinge helix. This residue is deleted in around 70% of all cystic fibrosis patient and results in a trafficking defect due to folding defects^58^. Small molecule correctors that stabilize the TMD can rescue these defects^59^. Similarly, our observation that hinge mutants that are misfolded in the context of the isolated BmrA NBD could be rescued in the presence of the TMD suggest that a destabilized NBD may generally require a stable TMD platform to enable proper folding. This highlights the value of studying isolated NBDs to derive mechanistic details of ABC transporters^60^, while also emphasizing the importance of the coupling helices for transporter stability.

While other ABC transporter classes lack the swapped domain topology of type IV transporters and the overall sequence conservation is low (**Fig. 4C**), some structural features persist throughout the superfamily. This includes a short helix near the C-terminal tip of the Walker A helix with a hydrophobic residue (position ϕ) followed by, in most cases, a positively charged residue (R^ICD2^), which points towards the respective intracellular domain (**Fig. 4B**). In some cases, such as in the type VI lipid transporter LptB_2_FG^61^ and the maltose transporter MalFGK_2_^62^ there is a positional switch in the sidechain properties at positions ϕ and R^ICD^ raising the question whether the flow of information on nucleotide and substrate binding is merged in a similar fashion throughout the whole ABC transporter family.

In conclusion, we identified a novel, interdomain transmission pathway in a type IV ABC transporter that functions as a dynamic bidirectional relay integrating multiple signals to ensure proper transporter function. This not only positions the hinge as a central regulator of interdomain communication in an important ABC transporter, but also provides a framework for studying similar relay elements in other complex biomacromolecular machines.

## Material and Methods

### Materials

All chemicals were purchased from Sigma-Aldrich, Roth and VWR unless otherwise stated. Reagents used include doxorubicin (Cayman Chemicals and Sigma), ^15^N-NH_4_Cl, ^13^C_6_-glucose (Eurisotop), 5-fluorotryptophane (BLD pharma) and ATTO oxa11 maleimide (ATTO-TEC). Lipids were purchased from Avanti Polar Lipids and Cayman Chemicals.

### Computational tools

Freely available computational tools were used to investigate the properties of BmrA constructs: Sequence alignment was carried out with Clustal Omega^70^ (https://www.ebi.ac.uk/jdispatcher/msa/clustalo) and sequence logos created using WebLogo 3^31^ (https://weblogo.threeplusone.com/). To create the WebLogo, 226 sequences were retrieved from the UniProtKB subdatabase (as of Nov. 27, 2024) which is part of the UniRef50_O06967 (50%) database. The latter corresponds to non-redondant protein sequences which show at least 50% sequence identity with BmrA. The sequences in the UniProtKB were either manually annotated and computationally analyzed (1 entry, UniProtKB reviewed (Swiss-Prot)) or only computationally curated from (227 entries, UniProtKB Unreviewed (TrEMBL)) to remove notably protein fragments. One entry was still a protein fragment (A0A7Y8S210_BACSP) and a second one contains one residue insertion between the Walker A and the Q-loop motif (A0A398D0Y7_9BACL). These two sequences were removed from our database and the remaining 226 sequences were used to make the WebLogo shown in Figure 1. NMR spectra were analyzed with Bruker TopSpin 4.0.8, CARA^71^ and CCP NMR^72^. HDX-MS data were analyzed with PLGS™ software (ProteinLynx Global SERVER 3.0.2 from Waters™), DynamX 3.0 software (Waters™), and Deuteros 2.0 software^73^.

### Cloning, expression, and purification of nucleotide binding domains (NBDs)

Synthetic genes coding for WT BmrA NBD (residues G331 – G589) from *Bacillus subtilis* and WT LmrA-NBD (residues D330 – Q590) from *Lactococcus lactis* cloned into pET-11a vector with a N-terminal His_6_-tag followed by a TEV cleavage site were obtained from GenScript (Piscataway Township, NJ, USA). Gene coding for WT MsbA-NBD (residues 323 – 582) was amplified from *E. coli* BL21(DE3) gold (Agilent Technologies) and cloned into pET-11a vector with a N-terminal His_6_-tag followed by a TEV cleavage site via Gibson assembly. Point mutations R389A, R389E, R389K, R389M, K380A, W413A, W413L, W413F, W413Y, R414A and R414K, C436S, N459W and S516C for BmrA-NBD, R397 and W421 for LmrA NBD and R391A, L415A and L415W for MsbA NBD were introduced via site directed mutagenesis (see **Supplementary Table S5** for primer sequences).

Transformed *E. coli* BL21 (DE3) Gold cells were growth at 37 °C in 1L of Lysogeny Broth (LB) media supplemented with 100 µg/mL ampicillin until an OD_600_ of 0.6 was reached. Immediately a final concentration of 1 mM IPTG was added and cells were grown at 21 °C overnight. Cells were harvested by centrifugation (5000xg, 15 min, 4 °C) and the resulting cell pellet was frozen in liquid nitrogen and stored at − 20 °C until further use. Purification followed our previously established protocol^30^, but with an additional final concentration of 2.5 mM ADP added during lysis for R389A, R389E, R389K and R389M constructs to prevent protein aggregation.

### Labeling of nucleotide binding domains (NBDs) for NMR and PET-FCS

Proteins were expressed and purified as described above with some modifications. ^1^H, ^15^N-labelled BmrA NBD WT, K380A, R389K, R389M, W413F, R414A and R414K for NMR spectroscopy were obtained by growing cells in M9 minimal medium^74^ supplemented with ^15^N-NH_4_Cl as the sole nitrogen source. ^19^F-tryptophane-labelled BmrA NBD WT, K380A, R389K, R389M and R414K were obtained by growing cells in defined medium^74^ supplemented with 50 mg/L 5-fluorotryptophane.

To obtain fluorescently modified NBD constructs for PET-FCS, Ni-NTA-column-bound single cysteine mutants were incubated for 1 h at room temperature with 10 mM TCEP. Afterwards, resin was washed with 10 column volumes (CV) washing buffer (50 mM Tris HCl pH 8, 500 mM NaCl) and incubated with 15-fold excess of ATTO oxa11 maleimide (ATTO-TEC) for 2 h at room temperature. To remove excess of fluorophore, the resin was washed with 10 CV of washing buffer (50 mM Tris HCl pH 8, 500 mM NaCl) and labelled protein was eluted and purified as previously described^30^.

### Cloning, expression and purification of membrane scaffold protein MSP1E3D1

Expression and purification of MSP1E3D1 was performed as previously described^75^. In brief, using the p1E3D1 plasmid (Addgene), MSP1E3D1 was expressed in *E. coli* BL21 cells. After harvest, bacteria were suspended in 50 mL of 40 mM Tris-HCl pH 7.4, 100 mM NaCl, 1 % (w/v) Triton X100, 0.5 mM EDTA, 1 mM PMSF. Two microliters of Benzonase (24 U/mL, Merck) were added and the bacteria were lysed with two passages at 18,000 psi through a microfluidizer 100 (Microfluidics IDEX Corp) and then centrifuged during 30 min. at 30,000 xg, 4°C. The supernatant was loaded onto a 0.5-mL Ni^2+^-NTA column (GE Healthcare) resin pre-equilibrated with 5 resin-volumes of 40 mM Tris-HCl pH 7.4, 100 mM NaCl, 1 % (w/v) Triton X100, 0.5 mM EDTA and 1 mM PMSF. The resin was then washed with 10 resin-volume with 3 different buffers: wash buffer 1 composed of 40 mM Tris-HCl pH 8.0, 300 mM NaCl and 1% (w/v) Triton X100; wash buffer 2 composed of 40 mM Tris-HCl pH 8.0, 300 mM NaCl, 50 mM sodium cholate and 20 mM Imidazole; wash buffer 3 composed of 40 mM Tris-HCl pH 8.0, 300 mM NaCl, 50 mM imidazole. MSP1E3D1 was eluted with 15 mL of 40 mM Tris-HCl pH 8.0, 300 mM NaCl and 500 mM Imidazole. TEV (2 mg/mL, with 1 mg TEV for 40 mg MSP1E3D1) was added to remove the His tag during dialysis (using a 12-14 kDa cutoff membrane) against 300 mL 40 mM Tris-HCl, pH 7.4, 100 mM NaCl and 0.5 mM EDTA for 3 hours and then against 700 mL of the same buffer, overnight at 4 °C. After dialysis, 20 mM imidazole was added and the solution loaded on a 0.5 mL Ni^2+^-NTA column equilibrated with 20 mM Tris-HCl pH 7.4 and 100 mM NaCl to remove uncleaved protein and the His-tagged TEV protease. MSP1E3D1 was concentrated spinning at 5,000 xg with a 10 kDa cutoff Amicon Ultra-15. The concentrated samples were frozen in liquid nitrogen and stored at −80 °C.

### Cloning and expression of full-length BmrA constructs

Vector pET23a (+) coding for BmrA full-length WT with a C-terminal His_6_ tag was obtained as previously described^8^. Point mutations for R389A, R389E, R389K, R389M, W413A, W413L, W413F, W413Y, R414A, R414K and K380A were introduced via site directed mutagenesis (see **Supplementary Table S5** for primer sequences). Transformed *E. coli* C41 (DE3) cells were growth overnight at 37 °C in a 100 mL of LB culture. The overnight preculture was then used to inoculate 1 L of 2xYT medium containing 100 μg/ml ampicillin to an optical density (*A*_600 nm_) of 0.05. The culture was then incubated at 37 °C until an OD_600_ of 0.6 was reached. ^19^F-tryptophane-labelled WT, W104F, W164F and W413Y BmrA full-length were obtained by growing cells in defined medium^74^ supplemented with 50 mg/L 5-fluorotryptophane. For expression induction, a final concentration of 700 µM IPTG was added and cells were grown at 25 °C for four hours. Afterwards, cells were harvested by centrifugation (5000 xg, 15 min, 4° C) and the resulting cell pellet was frozen in liquid nitrogen and stored at − 80 °C until further use.

### Inside – Out Vesicle (IOV) preparation

For membrane isolation, the frozen cell pellet was resuspended in lysis buffer (50 mM Tris/HCl pH 8, 5 mM MgCl_2_, 1 mM DTT) supplemented with 5 µg/mL of DNase I according to available protocols^8,76^. To disrupt the cells, the suspension was passed three times through a microfluidizer at 18,000 psi. The lysate was centrifuged at 15,000 g for 30 min at 4 °C to remove cell debris and insoluble proteins. After centrifugation, the supernatant containing the membranes was ultracentrifuged at 150,000 g for at least 1 h at 4 °C to harvest cell membranes. The supernatant was discarded and the pellet containing membranes was resuspended in resuspension buffer (50 mM Tris/HCl pH 8, 1.5 mM EDTA, 1 mM DTT). To reduce possible impurities in the membranes, a second ultracentrifugation step was performed at 150,000 xg for 1 h at 4 °C. The pellet from the second ultracentrifugation step was homogenized with a Dounce homogenizer in homogenization buffer (50 mM Tris/HCl pH 8, 1 mM EDTA, 300 mM sucrose) and aliquots were frozen in liquid nitrogen and stored at − 80 °C until further use.

For further analysis, total membrane protein was quantified in the IOVs using the Colorimetric bicinchoninic Acid (BCA) assay (Thermo Fisher Scientific, Waltham, USA) where absorption at 562 nm was measured to determine the protein concentration.

### Purification of full-length BmrA constructs

Full-length BmrA wildtype or its mutants were purified by solubilizing the membrane proteins, and diluting IOVs containing the overexpressed transporters to 2 mg/mL of total membrane protein using solubilization buffer (50 mM Tris/HCl pH 8, 50 mM NaCl, 10 mM imidazole, 1 mM DTT, 10% (v/v) glycerol and 1% (v/v) n-dodecyl–ß–D– maltopyranoside (DDM))^27^. The solution was incubated for 1 h at 4 °C and 200 rpm and then ultracentrifuged at 150,000 xg for 1 h at 4 °C to remove insoluble proteins. The clear supernatant was loaded onto a Ni-NTA column equilibrated as previously described^27^ or incubated with NiNTA beads previously equilibrated with equilibration buffer (50 mM Tris/HCl pH 8, 50 mM NaCl, 10% (v/v) glycerol, 10 mM imidazole pH 8, 0.0675% (v/v) DDM and 0.04% (v/v) Na–Cholate) for 1.5 h at 4 °C and 200 rpm. Afterwards, the mixture was transferred to a gravity flow column and washed with 5 CV of washing buffer (Tris/HCl pH 8, 500 mM NaCl and 10% (v/v) glycerol) followed by 20 cv of buffer wash A (50 mM Tris/HCl pH 8, NaCl 50 mM, 10% (v/v) glycerol, 25 mM imidazole, 0.0675% (v/v) DDM and 0.04% (v/v) Na–Cholate). BmrA constructs were eluted either using an imidazole gradient from 20 to 500 mM^27^ or with 6 CV of elution buffer (50 mM Tris/HCl, 50 mM NaCl, 10 % (v/v) glycerol, 250 mM Imidazole pH 8, 0.0675% (v/v) DDM and 0.04% (v/v) Na– Cholate). To reduce the imidazole concentration and to perform a buffer exchange, proteins were concentrated at 5,000 xg with a 100 kDa cutoff Amicon Ultra-15 and loaded onto a size exclusion column (Superdex200 Increase 10/300 GL (Cytiva)) previously equilibrated with SEC buffer (50 mM Hepes/KOH pH 8, 50 mM NaCl, 10% (v/v) glycerol, 0.035% (v/v) DDM and 0.03% (v/v) Na– Cholate) via an NGC chromatography system (Bio–Rad). The fractions containing pure protein were pooled and sample purity was verified by SDS–PAGE. Aliquots were frozen in liquid nitrogen and stored at − 80 °C until further use.

### Reconstitution in liposomes

Proteoliposomes were prepared as described in Orelle et al., 2003^77^. Briefly, a stock of 25 mg/mL *E. coli* lipid extract was prepared in autoclaved MilliQ water. 10 µL of 10% DDM were added to 40 µL of the lipid stock in a 2 mL reaction tube at room temperature. After 1 h at constant stirring, 50 µg of previously purified transporter in DDM/Na– Cholate were added and the final volume adjusted to 250 µL with reconstitution buffer (50 mM Hepes/KOH pH 8 and 50 mM NaCl). After 1 h incubation at room temperature and constant stirring, three subsequent additions of 20 mg of dried Bio–Beads SM2 were done every hour. Afterwards, the mixture was centrifugated at 6000xg and supernatant containing proteoliposomes was kept at 4 °C until further use.

### Reconstitution in nanodiscs

Membrane Scaffold Protein (MSP) nanodiscs were prepared following the protocol by Alvarez et al^75^. A stock of 25 mg/mL *E. coli* lipid extract was prepared in autoclaved MilliQ water. 9.3 µL of the lipid stock were mixed with 15 µL of 10% DDM in a 2 mL reaction tube at room temperature. After 1 h at constant stirring, 100 µg of purified transporter in DDM/Na–Cholate and 96 µg of purified MSP1E3D1 were added and the final volume adjusted to 250 µL with reconstitution buffer (50 mM Hepes/KOH pH 8 and 50 mM NaCl). After 1 h of incubation at room temperature and constant stirring, 170 mg of dried Bio–Beads SM2 were added and the mixture was incubated for 3 h at room temperature and constant stirring. To obtain the nanodisc reconstituted ABC transporter, the mixture was centrifuged at 6000 xg and the supernatant containing nanodiscs was kept at 4 °C until further use.

### Doxorubicin transport assay

The transport of doxorubicin by overexpressed BmrA transporters was studied in IOVs. The excitation and emission wavelengths were set at 480 and 590 nm, respectively. One hundred µg of IOVs containing BmrA wildtype or variants were diluted to 1 mL with transport buffer (50 mM Hepes/KOH pH 8, 8.5 mM NaCl, 4 mM phosphoenolpyruvate (PEP), 2 mM MgCl_2_ and 60 µg of pyruvate kinase) in a 10 mm quartz cuvette (Hellma Analytics, Müllheim) and placed in a fluorimeter (Horiba or Photon Technology International, Inc.). Measurements were carried out at 25 °C under constant stirring with spectral bandwidths of 2 and 4 nm for excitation and emission, respectively. Fluorescence was recorded for 125 s before 10 µM of doxorubicin was added and the fluorescence was recorded for an additional 125 s. After these 250 s, ATP was added to a final concentration of 2 mM and the measurement continued for another 350 s.

### ATPase activity

The ATPase activity of detergent-purified or reconstituted BmrA constructs was determined based on an ATP–regenerating system coupling ATP hydrolysis to NADH oxidation which can be monitored spectrophotometrically at 340 nm. Using previously published protocols with slight modifications^8,76^, NADH oxidation was monitored by measuring loss of absorbance at 340 nm in an UV–Vis spectrophotometer (SAFAS SP2000, Monaco) at 37 °C over 20 min. For each measurement, 3 µg of BmrA in detergent micelles or 1 µg of reconstituted transporter in nanodiscs or liposomes were diluted to 700 µL with ATPase buffer (50 mM Hepes/KOH pH 8, 10 mM MgCl_2_, 4 mM phosphoenolpyruvate, 32 µg/mL lactate dehydrogenase, 60 µg/mL pyruvate kinase and 0.3 mM NADH) in a 10 mm quartz cuvette (Hellma Analytics, Müllheim). For the protein in detergent micelles, DDM and Na–Cholate were added to final concentrations of 0.035% (v/v) and 0.03% (v/v), respectively.

### Analytical size-exclusion chromatography (SEC)

20 µM of purified ABC transporter constructs, either isolated NBDs in 50 mM BisTris pH 7, 50 mM NaCl or full-length constructs in 50 mM Hepes/KOH pH 8, 50 mM NaCl, 10% glycerol, 0.035% DDM and 0.03% Na–Cholate were used. Samples were injected on a Superdex200 Increase 10/300 GL (Cytiva) column via an NGC chromatography system (Bio–Rad).

### Circular dichroism (CD) spectroscopy

CD measurements were conducted on a Jasco J–1500 CD spectrometer (Jasco, Gross–Umstadt, Germany) with 1 mm quartz cuvettes. For isolated NBD constructs, 5 µM protein in 5 mM Tris pH 7 and 2.5 mM NaCl were used. For the whole transporters, 1 µM protein was used in 0.5 mM Hepes/KOH pH 8, 1.5 mM NaCl, 0.035% DDM and 0.03% Na–Cholate. Spectra were recorded at 25 °C in a spectral range between 190 and 260 nm with 1 nm scanning intervals, 1.00 nm bandwidth and 50 nm/min scanning speed. All spectra were obtained from the automatic averaging of five measurements.

### Thermal stability assays

Nano differential scanning fluorimetry using the Prometheus NT.48 instrument (Nanotemper technologies, DE) was used^27,78^. BmrA samples at 1 mg/mL were incubated for 15 min at room temperature in the absence of ligands or in the presence of 10 mM ATP, 10 mM MgCl_2_ and 1 mM sodium orthovanadate (V_i_). 10 µL of the sample was used to fill the capillaries. A temperature gradient of 1 °C/min from 20 to 95 °C was applied and fluorescence was recorded at 330 and 350 nm. The ratio of fluorescence intensities at 350/330 nm was used to determine the melting temperature (*T_m_*).

In addition, the melting temperatures (*T_m_*) of the protein constructs were determined with a fluorescence-based assay^79^. Purified protein (5-25 µg) in 50 mM BisTris pH 7, 50 mM NaCl were used. Proteins were measured in the absence of ligands or incubated with 10 mM ADP or ATP with and without 10 mM MgCl_2_. 2.5 µL of a 50X SYPRO Orange dye stock was added to each sample directly before measuring the melting temperature in a 96-well plate. Measurements were carried out on a QuantumStudio 1 Real–Time PCR System reader (Thermo Fisher) with a temperature increase of 0.05 /min. The fluorescence of SYPRO Orange was measured using for a SYBR GREEN–calibrated filter with an excitation filter of 470 +/– 15 nm and an emission of 520 +/– 15 nm.

For ABC transporters WT, K/A and R389X mutants, experiments were carried out with 10–25 µg of purified protein in Hepes/KOH pH 8, 50 mM NaCl, 10% (v/v) glycerol, 0.035% (v/v) DDM and 0.03% (v/v) Na–Cholate. The samples were measured in the absence of ligands or incubated with 10 mM ADP or ATP with 10 mM MgCl_2_. 2 µL of a 10X GloMelt fluorescent dye stock was added to each sample directly before measurement in a 96-well plate. The melting temperatures were determined with a QuantumStudio 1 Real–Time PCR System reader (Thermo Fisher) with a temperature increase of 0.05 °C /min. The fluorescence of GloMelt was measured using the filter calibrated for SYBR GREEN with an excitation filter of 470 +/– 15 nm and an emission of 520 +/– 15 nm.

### Solution NMR spectroscopy

For NMR experiments, samples were concentrated to 200 – 400 µM before addition of 10 mM ADP or ATP and 10% v/v D_2_O and 0.15 mM DSS (final concentrations). TROSY – based ^1^H, ^15^N-HSQC NMR spectra of isotope labeled NBDs in 50 mM BisTris pH 7, 50 mM NaCl were recorded at 298 K on a Bruker AVANCE 600 or Bruker Neo 800 MHz spectrometer equipped with cryogenic triple resonance probes (Bruker GmbH, Karlsruhe, Germany) using standard NMR pulse sequenced implemented in Bruker Topspin library. All spectra were processed using Bruker TOPSPIN 4.0.8 and analyzed using CCP NMR analysis^72^.

The average ^1^H and ^15^N weighted chemical shift perturbations (CSP) observed in ^1^H, ^15^N HSQC spectra can be calculated according to equation 1^80^:

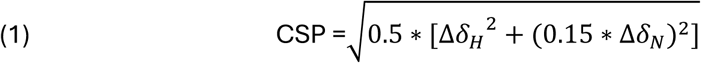

Here, Δ*δ*_*H*_ is the ^1^H chemical shift difference, Δ*δ*_*N*_ is the ^15^N chemical shift difference, and CSP is the average ^1^H and ^15^N weighted chemical shift difference in ppm.

### 19F NMR

^19^F-NMR spectra of 5-fluorotryptophane labeled BmrA NBD and full-length constructs were recorded at 298 K on a Bruker AVANCE III 600 MHz spectrometer equipped with a Prodigy TCI or a CP2.1 QCI cryoprobe (Bruker GmbH, Karlsruhe, Germany) using standard NMR pulse sequences implemented in Bruker Topspin library. For isolated NBDs all measurements were carried out in 50 mM BisTris pH 7, 50 mM NaCl. Samples were concentrated to 100 – 250 µM before addition of 10 mM ADP and 10% v/v D_2_O and 0.15 mM DSS (final concentrations). In the case of 5-fluorotryptophane labeled BmrA full-length constructs, all measurements were carried out in Hepes/KOH pH 8, 50 mM NaCl, 10% (v/v) glycerol, 0.035% (v/v) DDM and 0.03% (v/v) Na– Cholate. Samples were concentrated to 100 – 300 µM before addition of 10 mM ADP, 10 mM ATP or 15 mM reserpine and 10% v/v D_2_O and 0.15 mM DSS (final concentrations). All spectra were processed using Bruker TOPSPIN 4.0.8 and deconvoluted using pure Lorenztian-line-shape fitting.

### Photoinduced electron transfer fluorescence correlation spectroscopy (PET-FCS) experiment

PET-FCS was performed using a custom-built confocal fluorescence microscope setup described elsewhere^81^. Fluorescently modified NBD constructs were diluted to 1 nM final concentration in 50 mM phosphate buffer pH 7.0 with the solution ionic strength adjusted to 200 mM using potassium chloride. 0.3 mg/ml Protease-free bovine serum albumin and 0.05% Tween-20 were applied as solution additives to suppress sample/glass-surface interactions. The buffer was filtered through a 0.2 µm syringe filter. To study the effect of nucleotides, 1 nM protein samples were incubated with 10 mM ADP or ATP directly prior to measurement and then transferred onto a microscope slide and covered by a cover slip. The sample temperature was adjusted to 25 °C using a custom-built objective heater. The total measurement time was 15 minutes for each sample.

### PET-FCS data analysis

Autocorrelation functions were analyzed by fitting a model for a two-dimensional diffusion of a globule containing a sum of *n* single-exponential relaxations^38,82^:

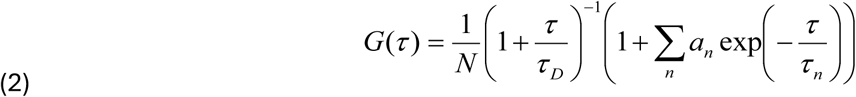

*t* is the lag time, *N* is the average number of molecules in the detection volume, *τ_D_* is the experimental diffusion time constant, *a*_n_ and *τ*ν are the observed amplitude and time constant of the n^th^ relaxation. The application of a model for diffusion in two dimensions was of sufficient accuracy because the two horizontal dimensions (*x*, *y*) of the detection focus were much smaller than the lateral dimension (*z*) in the applied setup.

### Hydrogen Deuterium Exchange coupled to Mass Spectrometry

HDX-MS experiments were performed using a Synapt G2-Si mass spectrometer coupled to a NanoAcquity UPLC M-Class System with HDX Technology (Waters™). All the reactions were carried out manually, as previously described^26^. Deuteration labeling was initiated by diluting 5 μL of 20 μM BmrA W413F or 5 μL of 15 μM BmrA WT reconstituted in nanodiscs in 95 μL D_2_O labeling buffer (5 mM Hepes pD 8.0, 50 mM NaCl). For ADP/Vi trapping, the labeling buffer additionally contained 10 mM ATP, 10 mM MgCl_2_, 1 mM Vi. For the doxorubicin-bound condition, the labeling buffer additionally contained 100 μM doxorubicin. Prior to labeling, the samples were incubated with the respective ligands for 15 min at 20 °C. Samples were labeled for 2, 5, 15 and 30 min at 20 °C. Subsequently, the reactions were quenched by adding 22 μL of ice-cold quenching buffer (0.5 M glycine, 8 M guanidine-HCl pH 2.2, 0.035% DDM and 0.03% sodium cholate) to 100 μL of labeled sample, in ice bath. After 1 min, the 122-μL quenched sample was added into a microtube containing 200 μg of activated zirconium magnetic beads (MagReSyn Zr-IMAC from Resyn Biosciences, USA) to remove the phospholipids ^83^. After 1 min, magnetic beads were removed, and the supernatant was injected immediately into a 100-μL loop. Labeled proteins were then subjected to on-line digestion at 15 °C using a pepsin column (Waters Enzymate™ BEH Pepsin Column 300 Å, 5 μm, 2.1 x 30 mm). The resulting peptides were trapped and desalted for 3 min on a C4 pre-column (Waters ACQUITY UPLC Protein BEH C4 VanGuard pre-column 300 Å, 1.7 μm, 2.1 x 5 mm, 10K - 500K) before separating them with a C4 column (Waters ACQUITY UPLC Protein BEH C4 Column 300 Å, 1.7 μm, 1 × 100 mm) using 0.2% formic acid and a 5-40 % linear acetonitrile gradient in 15 min and then 4 alternative cycles of 5% and 95% until 25 min. The valve position was adjusted to divert the sample after 14 min of each run from C4 column to waste, to avoid a contamination of the mass spectrometer with detergent. At least two full kinetics were run for each condition, directly subsequently. Blanks (equilibration buffer: 5 mM Hepes pH 8.0, 50 mM NaCl) were injected after each sample injection and pepsin column washed during each run with pepsin wash (1.5 M guanidine-HCl, 4% acetonitrile, 0.8% formic acid pH 2.5) to minimize the carryover. Electrospray ionization mass spectra were acquired in positive mode in the m/z range of 50−2000 and with a scan time of 0.3 s. For the identification of non-deuterated peptides, data was collected in MSE mode and the resulting peptides were identified using PLGS™ software (ProteinLynx Global SERVER 3.0.2 from Waters™). Peptides were then filtered in DynamX 3.0 software (Waters™), with the following parameters: minimum intensity of 1000, minimum products per amino acid of 0.3 and file threshold of 2 out of 8 to 11. After manual curation, Deuteros 2.0 software was used for data analysis, visualization and statistical treatments for differential HDX-MS. The significance of identified changes of deuteration was evaluated using peptide significance statistical tests^73^ using p-value threshold of 0,05. Unsignificant peptides were then manually removed in DynamX 3.0 software (Waters™). Data were finally integrated in PDB files using HDX-viewer tool^84^ for 3D visualization. Figures were prepared using the PyMOL Molecular Graphics System (Version 3.0 Schrödinger, LLC^85^). The mass spectrometry data have been deposited to the ProteomeXchange Consortium via the PRIDE partner repository with the dataset identifier PXD027447 for BmrA WT in the apo and ADP*Vi states and PXD058013 for the W413F mutant in the respective states as well as WT and mutant bound to doxorubicin.

### Data availability

The backbone NMR assignments of *L. lactis* LmrA^29^, *B. subtilis* BmrA^30^ and *E. coli* MsbA have been deposited to the BMRB under the accession numbers 51156, 17660, and 52626, respectively. The mass spectrometry data have been deposited to the ProteomeXchange Consortium via the PRIDE^86^ partner repository with the dataset identifier PXD058013.

## Acknowledgements

CO and UAH would like to dedicate this article to the memory of Amy L. Davidson, who first brought us together, but then left way too soon. We thank Elise Kaplan, Mai Anh Tran, Jan Overbeck and Christoph Wiedemann for fruitful discussions and technical support. VHPC acknowledges a DAAD-CONACYT PhD fellowship and a PROCOPE mobility 2020 fellowship by the Department for Science and Technology of the Embassy of France in Germany. DRS acknowledges a PhD fellowship, and a research stay fellowship by the Hans Böckler Stiftung. This work was supported by the Deutsche Forschungsgemeinschaft, Project ID 421388231 (to HN and UAH), and the Cluster of Excellence “Balance of the Microverse” (EXC 2051—Project-ID 390713860) (to UAH) as well as by the Agence Nationale de la Recherche grant N° ANR-19-CE11-0023-01 (to CO) including a PhD fellowship for MDC. We thank the Fondation pour la Recherche Médicale (FRM) for supporting the 6-months PhD extension for MDC (Funding number FDT202204015047). UAH acknowledges an instrumentation grant for a high-field NMR spectrometer by the REACT-EU EFRE Thuringia (Recovery assistance for cohesion and the territories of Europe, ERDF, Thuringia) initiative of the European Union. CO and J-MJ acknowledge the Protein Science Facility of SFR Biosciences (Universite Claude Bernard Lyon 1, CNRS UAR3444, Inserm US8, ENS de Lyon) for the use of the circular dichroism equipment.

