## Supplementary Information for "Bidirectional communication between nucleotide and substrate binding sites in a type IV multidrug ABC transporter"

**Supplementary Tables**

**Supplementary Table 1: Thermal stability of BmrA, LmrA and MsbA NBD communication hinge mutants.** Melting temperatures ($T_m$) of ABC transporter NBDs were determined without nucleotides (apo) and in the presence of 10mM ATP or ADP using a SYPRO Orange fluorescence assay. Results are shown as mean ± standard deviation. All measurements were performed with two biological replicates.

| Protein construct | | $T_{m,\ apo}$ (ºC) | $T_{m,\ MgADP}$ (ºC) | $T_{m,\ ATP}$ (ºC) | $T_{m,\ MgATP}$ (ºC) |
|---|---|---|---|---|---|
| BmrA NBD | WT | 41.1 ± 0.2 | 45.8 ± 0.5 | 44.0 ± 0.1 | 44.3 ± 0.1 |
| | K380A | 41.0 ± 0.1 | 40.5 ± 0.1 | 40.3 ± 0.1 | 40.4 ± 0.2 |
| | R389A | 33.8 ± 0.5 | 40.6 ± 0.4 | 39.6 ± 0.3 | 39.9 ± 0.3 |
| | R389E | 33.6 ± 0.1 | 37.5 ± 0.2 | 37.1 ± 0.1 | 37.4 ± 0.3 |
| | R389K | 34.9 ± 0.3 | 41.0 ± 0.3 | 39.3 ± 0.1 | 39.5 ± 0.2 |
| | R389M | 36.6 ± 0.3 | 40.5± 0.1 | 39.8 ± 0.1 | 39.9 ± 0.4 |
| | W413A | Non purifiable protein | | | |
| | W413L | Non purifiable protein | | | |
| | W413F | 45.9 ± 0.3 | 51.2 ± 0.4 | 51.0 ± 0.2 | 50.1 ± 0.3 |
| | W413Y | 36.8 ± 0.4 | 46.7 ± 1.0 | 47.0 ± 0.3 | 44.2± 0.4 |
| | R414A | 39.0 ± 0.1 | 46.8 ± 0.1 | 43.7 ± 0.2 | 44.5 ± 0.3 |
| | R414K | 39.5 ± 0.1 | 46.2 ± 0.2 | 43.7 ± 0.1 | 45.1 ± 0.4 |
| LmrA NBD | WT | 41.2 ± 1.7 | 46.1 ± 0.1 | 46.8 ± 0.9 | 45.7 ± 0.1 |
| | R397A | 34.9 ± 0.8 | 33.9 ± 1.5 | 33.5 ± 1.3 | 35.6 ± 2.3 |
| | W421A | 32.3 ± 0.1 | 33.1 ± 1.6 | 31.8 ± 0.6 | 33.6 ± 0.1 |
| MsbA NBD | WT | 47.2 ± 0.1 | 51.0 ± 0.1 | 52.1 ± 1.1 | 50.0 ± 0.3 |
| | R391A | 32.0 ± 0.5 | 44.8 ± 0.4 | 45.1 ± 0.1 | 43.8 ± 2.5 |
| | L415A | 37.6 ± 0.9 | 45.7 ± 0.1 | 44.4 ± 0.3 | 44.7 ± 0.5 |
| | L415W | 44.2 ± 0.6 | 49.1 ± 0.5 | 48.3 ± 0.5 | 47.4 ± 0.6 |

**Supplementary Table 2: Dissociation constants ($K_d$) of BmrA NBD mutants for ADP derived from chemical shift perturbation assays.**

| Construct | $K_{d\,ADP}$ (µM) |
|---|---|
| BmrA NBD WT | 214.1 ± 42.9 |
| BmrA NBD K380A | 14410 ± 3700 |
| BmrA NBD R389K | 220.8 ± 105.0 |
| BmrA NBD R389M | 2887.8 ± 1461.3 |
| BmrA NBD W413F | 220.7 ± 43.3 |
| BmrA NBD R414A | 575.1 ±43.5 |
| BmrA NBD R414K | 265.6 ± 96.5 |

**Supplementary Table 3: Effect of hinge mutants on the thermal stability of full-length BmrA.** Melting temperatures ($T_m$) of protein in DDM/Cholate detergent micelles were determined using nanoDSF (R414X, W413X) or the fluorescent GloMelt™assay (R398X and $^{19}$F-5Trp-labeled samples) in the apo state (nucleotide free) and in presence of MgATPV$_i$ to stabilize the nucleotide bound state. Results are shown as mean ± standard deviation from two biological replicates measured three times or, in the case of the $^{19}$F-labeled samples, from one biological replicate measured three times.

| **Full-length BmrA construct** | ***Tm*** $_{apo}$ **(ºC)** | ***Tm*** $_{MgATPVi}$ **(ºC)** |
|---|---|---|
| **WT** | 43.6 ± 1.0 | 51.2 ± 0.3 |
| **K380A** | 44.0 ± 0.3 | 44.9 ± 0.1 |
| **R389A** | 38.7 ± 0.3 | 43.9 ± 3.0 |
| **R389E** | 39.5 ± 1.0 | 43.6 ± 0.4 |
| **R389K** | 40.1 ± 0.1 | 43.7 ± 0.8 |
| **R389M** | 38.5 ± 1.2 | 48.1 ± 1.7 |
| **W413A** | 39.6 ± 1.7 | 42.2 ± 1 |
| **W413L** | 34.7 ± 0.4 | 49.8 ± 0.5 |
| **W413F** | 36.5 ± 0.3 | 52.0 ± 0.4 |
| **W413Y** | 37.1 ± 0.1 | 50.7 ± 0.6 |
| **R414A** | 37.4 ± 0.0 | 38.0 ± 0.0 |
| **R414K** | 39.3 ± 0.1 | 40.6 ± 0.1 |
| **$^{19}$F–5W WT** | 42.6 ± 1.7 | 56.3 ± 2.4 |
| **$^{19}$F–5W W104F** | 37.2 ± 0.8 | 57.5 ± 0.3 |
| **$^{19}$F–5W W164F** | 43.0 ± 1.8 | 61.5 ± 1.2 |
| **$^{19}$F–5W W413Y** | 45.9 ± 2.2 | 58.4 ± 1.1 |

**Supplementary Table 4: Fitting parameters for PET-FCS.** Kinetic parameters obtained for the BmrA NBD single (S516C), double (N459W-S516C) and the triple (N459W-S516C-W413F) mutants in apo and after incubation with ADP or ATP.

| | $a_1$ | $\tau_1$ (μs) | $a_2$ | $\tau_2$ (μs) | $a_3$ | $\tau_3$ (μs) |
|---|---|---|---|---|---|---|
| **S561C** | 0.05±0.01 | 8±2 | -- | -- | -- | -- |
| **S516C-N459W** | 0.19±0.03 | 80±3 | 0.62±0.06 | 2.5±0.4 | 0.72±0.06 | 0.24±0.04 |
| **S516C-N459W-W413F** | 0.19±0.01 | 39±5 | -- | -- | 0.18±0.01 | 0.57±0.01 |
| **S516C-N459W +ADP** | 0.35±0.03 | ±16 | 0.56±0.04 | 1.5±0.2 | 0.42±0.04 | 0.13±0.03 |
| **S516C-N459W +ATP** | 0.22±0.08 | 253±16 | 0.51±0.06 | 7±2 | 0.95±0.08 | 0.30±0.04 |
| **S516C-N459W-W413F +ADP** | 0.18±0.01 | 104±12 | 0.18±0.01 | 2.7±0.2 | -- | -- |
| **S516C-N459W-W413F +ATP** | 0.19±0.02 | 177±40 | 0.20±0.01 | 5.7±0.7 | -- | -- |

$a_n$ and $\tau_n$ are the observed amplitudes and corresponding time constants. Errors are s.e. of fits to the data.

**Supplementary Table 5: Sequences of primers used to obtain hinge and tryptophan mutants**

| Primer name | DNA Sequence |
|---|---|
| BmrA NBD K380A forward | 5' – GAGCGGTGGCGGTGCGACCACCCTGTTC – 3' |
| BmrA NBD K380A reverse | 5' – GTTTGAACAGGGTGGTCGCACCGCCACCGC – 3' |
| BmrA NBD R389A forward | 5' – CTGTTCAAACTGCTGGAGGCGTTTTATAGCCCGAC – 3' |
| BmrA NBD R389A reverse | 5' – GCGGTCGGGCTATAAAACGCCTCCAGCAG - 3' |
| BmrA NBD R389E forward | 5' – CCTGTTCAAACTGCTGGAGGAATTTTATAGCC – 3' |
| BmrA NBD R389E reverse | 5' – GTCGGGCTATAAAATTCCTCCAGCAGTTTG – 3' |
| BmrA NBD R389K forward | 5' – CCTGTTCAAACTGCTGGAGAAATTTTATAGCC – 3' |
| BmrA NBD R389K reverse | 5' – GTCGGGCTATAAAATTTCTCCAGCAGTTTG – 3' |
| BmrA NBD R389M forward | 5' – CCTGTTCAAACTGCTGGAGATGTTTTATAGCC – 3' |
| BmrA NBD R389M reverse | 5' – GTCGGGCTATAAAACATCTCCAGCAGTTTG – 3' |
| BmrA NBD W413A forward | 5' – CTACAGCCTGGAGAGCGCGCGTGAACAC – 3' |
| BmrA NBD W413A reverse | 5' – CATAACCAATGTGTTCACGCGCGCTCTCCAG – 3' |
| BmrA NBD W413L forward | 5' – GATACCTACAGCCTGGAGAGCCTGCGTGAACAC – 3' |
| BmrA NBD W413L reverse | 5' – GTATTGGTTACACTTCACGCAGGCTCTCCAGGCTG – 3' |
| BmrA NBD W413F forward | 5' – GATACCTACAGCCTGGAGAGCTTTCGTGAACAC – 3' |
| BmrA NBD W413F reverse | 5' – CATAACCAATGTGTTCACGAAAGCTCTCCAGGCTG – 3' |
| BmrA NBD W413Y forward | 5' – CCTACAGCCTGGAGAGCTATCGTGAAC – 3' |
| BmrA NBD W413Y reverse | 5' – CATAACCAATGTGTTCACGATAGCTCTCCAG – 3' |
| BmrA NBD R414A forward | 5' – CTACAGCCTGGAGAGCTGGGCGGAACACATTG – 3' |
| BmrA NBD R414A reverse | 5' – CACCATAACCAATGTGTTCCGCCCAGCTCTC – 3' |
| BmrA NBD R414K forward | 5' – CTACAGCCTGGAGAGCTGGAAGGAACACATTG – 3' |
| BmrA NBD R414K reverse | 5' – CACCATAACCAATGTGTTCCTTCCAGCTCTC – 3' |
| LmrA NBD W421A forward | 5' – GATAGCGTGAGCCTGGAGAACGCGCGTAGCCAG – 3' |
| LmrA NBD W421A reverse | 5' – CTAACGAAGCCGATCTGGCTACGCGCGTTCTCCAG – 3' |
| LmrA NBD R397A forward | 5' – GCACCATTTTCAGCCTGCTGGAGGCGTTTTATCAG – 3' |
| LmrA NBD R397A reverse | 5' – CGCGGTCGGCTGATAAAACGCCTCCAGCAG – 3' |
| MsbA NBD R391A forward | 5' – CCATCGCCAGCCTGATCACGGCTTTTTACG – 3' |
| MsbA NBD R391A reverse | 5' – GCCTTCATCAATATCGTAAAAAGCCGTGATC – 3' |
| MsbA NBD L415A forward | 5' – GAGTATACCCTGGCGTCGGCACGTAAC – 3' |
| MsbA NBD L415A reverse | 5' – CAGAGCAACCTGGTTACGTGCCGACGC – 3' |
| MsbA NBD L415W forward | 5' – CGCGAGTATACCCTGGCGTCGTGGCGTAACC – 3' |
| MsbA NBD L415W reverse | 5' – CACCAGAGCAACCTGGTTACGCCACGACGC – 3' |

**Table S5 (continued): Sequences of primers used to obtain hinge and tryptophan mutants**

| | |
|---|---|
| BmrA NBD C436C forward | 5' – CCATTCGTGAGAACATCAGCTACGGTCTGG – 3' |
| BmrA NBD C436C reverse | 5' – CACGTTCCAGACCGTAGCTGATGTTCTC – 3' |
| BmrA NBD N459W forward | 5' – CGGAGATGGCGTATGCGCTGTGGTTCATTAAAG – 3' |
| BmrA NBD N459W reverse | 5' – GTTCGGCAGTTCTTTAATGAACCACAGCGCATAC – 3' |
| BmrA NBD S516C forward | 5' – CTGGATAGCCAGAGCGAAAAGTGCGTTCAGCAAG – 3' |
| BmrA NBD S516C reverse | 5' – CTTCCAGCGCTTGCTGAACGCACTTTTCGCTC – 3' |
| BmrA full-length K380A forward | 5' – GCGGCGGGGGAGCGACGACGCTGTTTAAG – 3' |
| BmrA full-length K380A reverse | 5' – CTTAAACAGCGTCGTCGCTCCCCCGCCG – 3' |
| BmrA full-length R389A forward | 5' – CGCTGTTTAAGCTGCTTGAAGCGTTTTATTCTCCG – 3' |
| BmrA full-length R389A reverse | 5' – CTGCAGTCGGAGAATAAAACGCTTCAAGCAGC – 3' |
| BmrA full-length R389E forward | 5' – CGCTGTTTAAGCTGCTTGAAGAATTTTATTCTCCG – 3' |
| BmrA full-length R389E reverse | 5' – CTGCAGTCGGAGAATAAAATTCTTCAAGCAGC – 3' |
| BmrA full-length R389K forward | 5' – CGCTGTTTAAGCTGCTTGAAAAATTTTATTCTCCG – 3' |
| BmrA full-length R389K reverse | 5' – CTGCAGTCGGAGAATAAAATTTTTCAAGCAGC – 3' |
| BmrA full-length R389M forward | 5' – CGCTGTTTAAGCTGCTTGAAATGTTTTATTCTCCG – 3' |
| BmrA full-length R389M reverse | 5' – CTGCAGTCGGAGAATAAAACATTTCAAGCAGC – 3' |
| BmrA full-length W413A forward* | 5' –ACCCGATATGCTCCCTCGCCGACTCGAGCGAGTAAGTAT– 3' |
| BmrA full-length W413L forward* | 5' –ACCCGATATGCTCCCTCAACGACTCGAGCGAGTAAGTAT– 3' |
| BmrA full-length W413F forward* | 5' –ACCCGATATGCTCCCTAAACGACTCGAGCGAGTAAGTAT– 3' |
| BmrA full-length W413Y forward* | 5' –ACCCGATATGCTCCCTATACGACTCGAGCGAGTAAGTAT– 3' |
| BmrA full-length R414A forward | 5' –ACTCGCTTGAATCGTGGGCGGAGCATATCGGGTATG– 3' |
| BmrA full-length R414A reverse | 5' –CATACCCGATATGCTCCGCCCACGATTCAAGCGAGT– 3' |
| BmrA full-length R414K forward | 5' –CTCGCTTGAATCGTGGAAGGAGCATATCGGGTATG– 3' |
| BmrA full-length R414K reverse | 5' –CATACCCGATATGCTCCTTCCACGATTCAAGCGAG– 3' |
| BmrA full-length W104F forward | 5' –CCGGGCTGCGGGAGTTATTATTTAAGAAATT– 3' |
| BmrA full-length W104F reverse | 5' –GGCAGCTTAATTAATTTCTTAAATAATAACTCC– 3' |
| BmrA full-length W164F forward | 5' –CAATCTTGTTTATTATGAACTTTAAGCTGACAC– 3' |
| BmrA full-length W164F reverse | 5' –CTAACACAAGCAGTGTCAGCTTAAAGTTCATAAT– 3' |

* Only one primer was used following the same cloning technique as in Orelle et al., 2003.

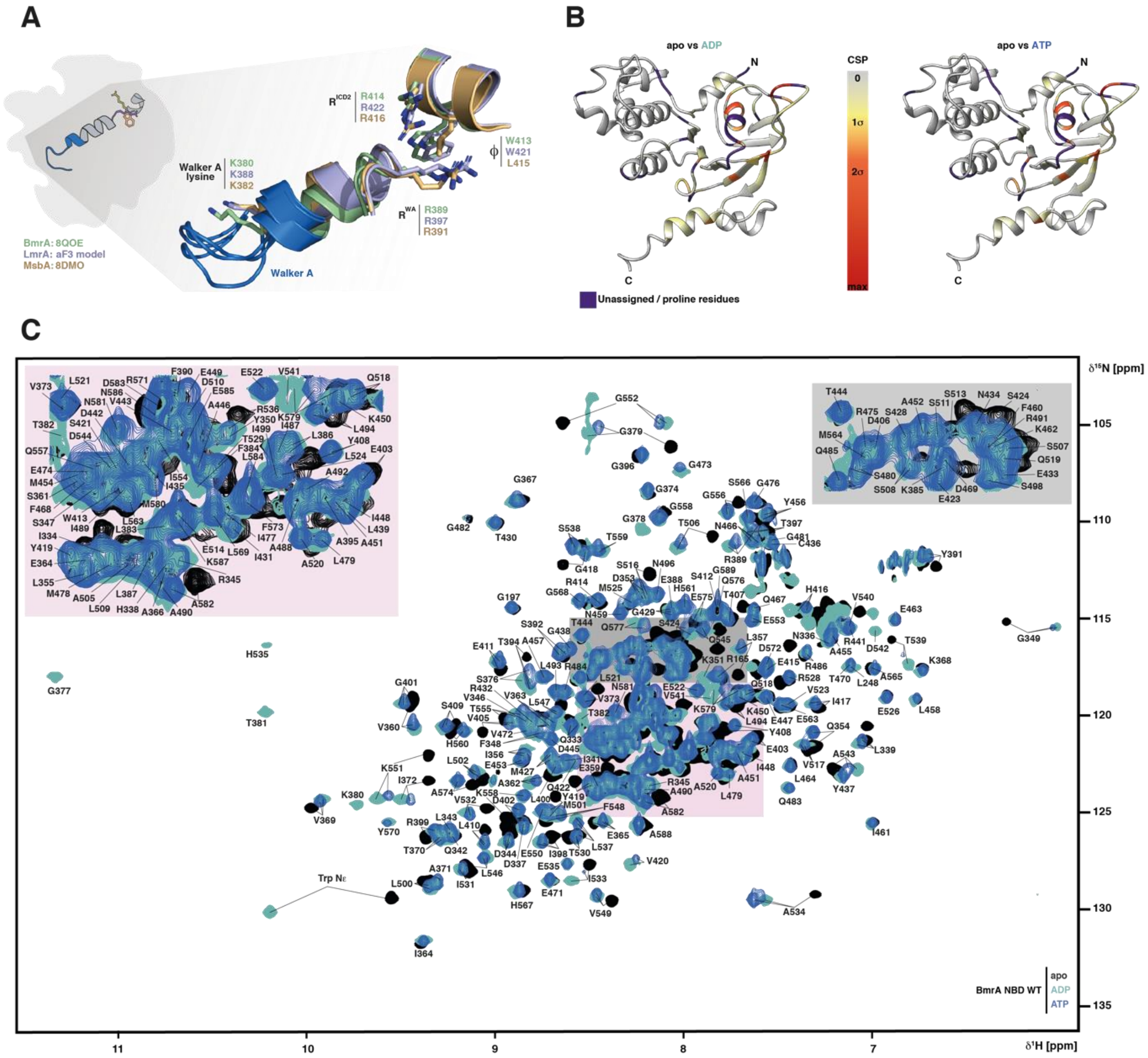


**Supplementary Figure 1: Structural conservation of hinge residues and nucleotide binding-induced chemical shift changes in the BmrA NBD.** (**A**) Structural overlay of the Walker A helix and hinge region in the NBD from *B. subtilis* BmrA (green, PDB ID: 8QOE (Di Cesare et al., 2024)), *L. lactis* LmrA (blue, AlphaFold3 model), and *E. coli* MsbA (gold, PDB ID: 8DMO (Lyu et al., 2022)) in the inward facing conformation. The conserved Walker A motif is shown in blue, the conserved Walker A lysine residue and the hinge residues are shown as coloured sticks. (**B**) Chemical shift perturbations (CSP) induced by the addition of 10 mM ADP (left) or ATP (right) to $^{15}$N-labeled BmrA NBD WT mapped onto the cryoEM structure of BmrA (PDB: 6R81 (Chaptal et al., 2022). Proline and other residues without amide backbone assignment in either state are colored in purple. CSP 1σ, 2σ and max values correspond to 0.01, 0.02 and 0.09 ppm respectively. (**C**) 2D–$^{1}$H,$^{15}$N–TROSY–HSQC spectrum of $^{2}$H, $^{15}$N–labeled BmrA–NBD in the apo state (black), with 10 mM ADP (teal) or 10 mM ATP (blue) were recorded at 298 K on a 600 MHz spectrometer equipped with a cryogenic triple resonance probe (Bruker GmbH, Karlsruhe, Germany), using samples with a concentration of 250 μM. To obtain the assignments of the apo and the ATP-bound states, we traced the chemical shift perturbations using our previously published backbone amide resonance assignment for BmrA bound to ADP (Pérez Carrillo et al., 2022) (BMRB entry 51156).

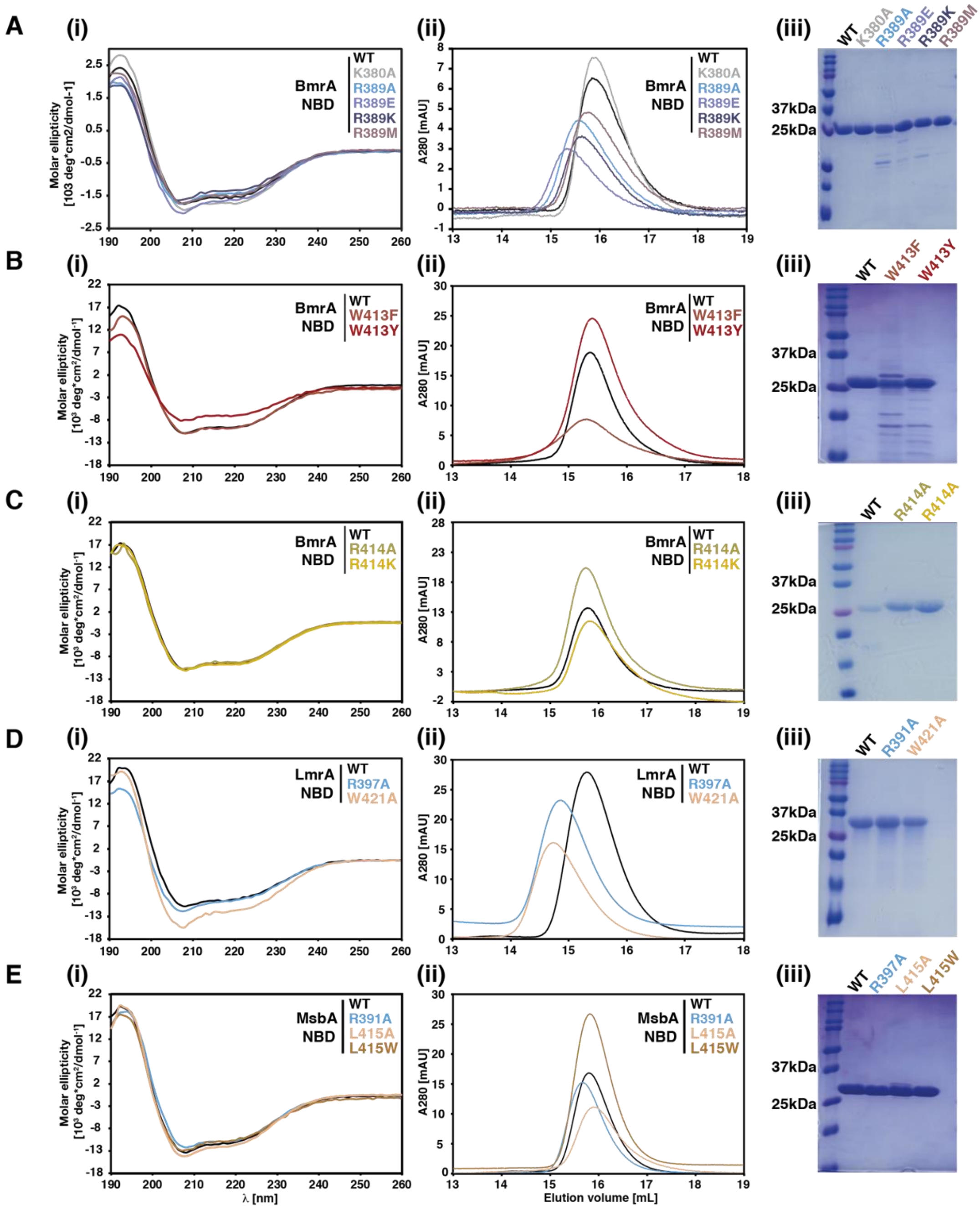


**Supplementary Figure 2: Structural integrity of the isolated NBDs from type IV ABC transporters containing hinge mutations. (A, B, C)** Circular dichroism spectra (i), Size exclusion chromatography profiles (ii) and SDS-PAGE (iii) of BmrA NBD with a mutation in residue $R^{WA}$ (A), ϕ (B) or $R^{ICD2}$(C), respectively. For reference, the BmrA NBD WT is included in (i-iii), and the Walker A K380A mutant in (i). (**D**) Circular dichroism spectra (i), Size exclusion chromatography profiles (ii) and SDS-PAGE (iii) of LmrA NBD WT and hinge mutants. (**E**) Circular dichroism spectra (i), size exclusion chromatography profiles (ii) and SDS-PAGE (iii) of MsbA NBD WT and hinge mutants.

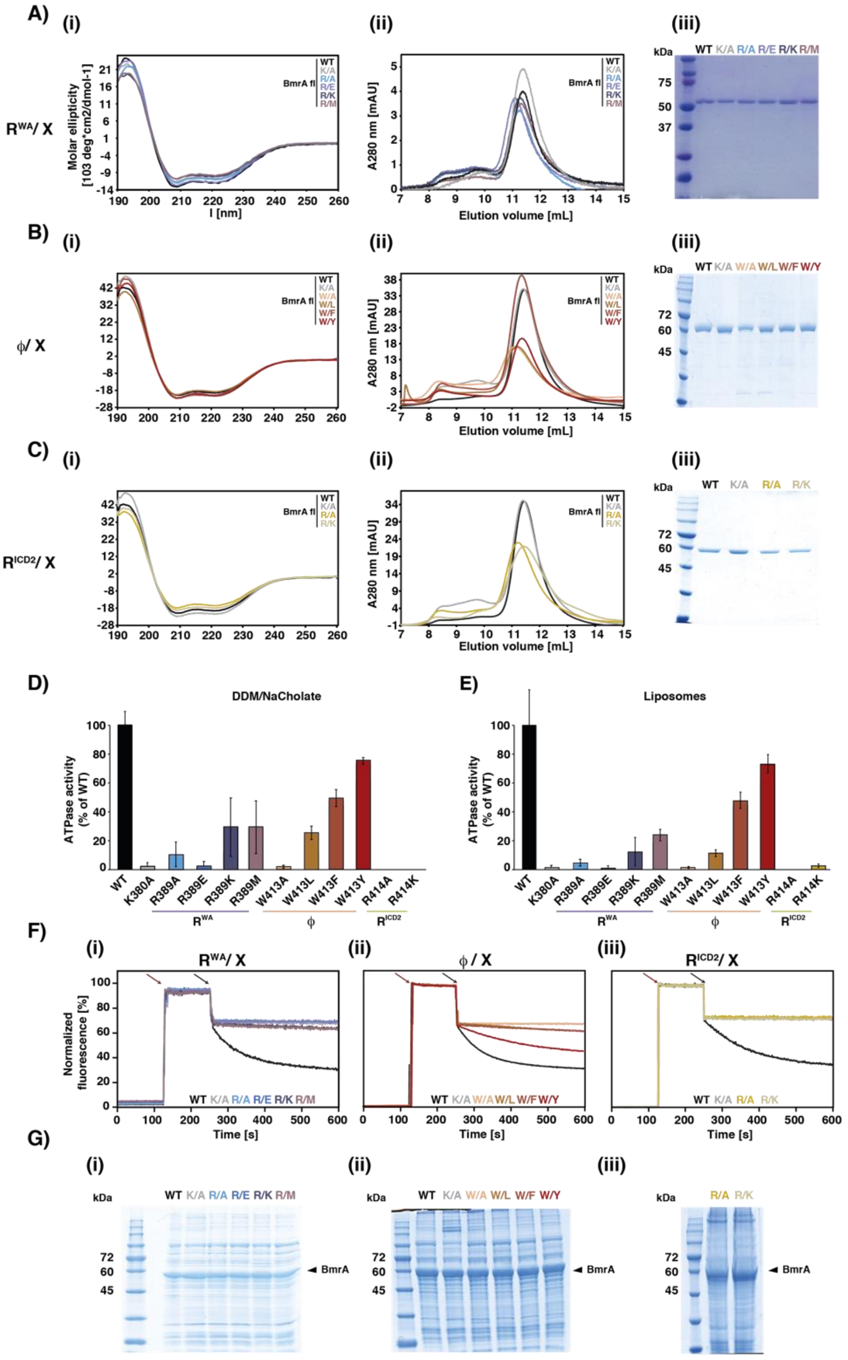

**Supplementary Figure 3: Role of hinge mutants for BmrA structural integrity and function.** (**A, B, C**) Circular dichroism spectra (i), size exclusion chromatography (ii) and SDS-PAGE of purified BmrA WT, Walker-A K380A, BmrA $R^{WA}$ (R398X), $\phi$ (W413X) and $R^{ICD2}$ (R414X) variants in DDM/cholate micelles. In (iii), 2.5 µg (A, B) or 1.0 µg (C) purified protein per lane were loaded (**D, E**) ATPase activity of BmrA variants in detergent micelles (DDM/NaCholate) (D) and reconstituted in liposomes prepared from *E. coli* polar lipid extract (E). All values were normalized to the WT activity. (**F**) Fluorescence-based doxorubicin transport assay with inside-out vesicles (IOVs) prepared from *E. coli* cells overexpressing *B. subtilis* BmrA (i) $R^{WA}$ (R398X), (ii) $\phi$ (W413X) and (iii) $R^{ICD2}$ (R414X) residue variants (see **Fig. 1E** for a quantitative analysis). Red arrows at 125 s and black arrows at 250 s indicate addition of the fluorescent substrate doxorubicin and ATP, respectively. (**G**) All variants were expressed similarly as seen by SDS-PAGE of IOVs. The expected running height of BmrA is indicated by a black arrow.

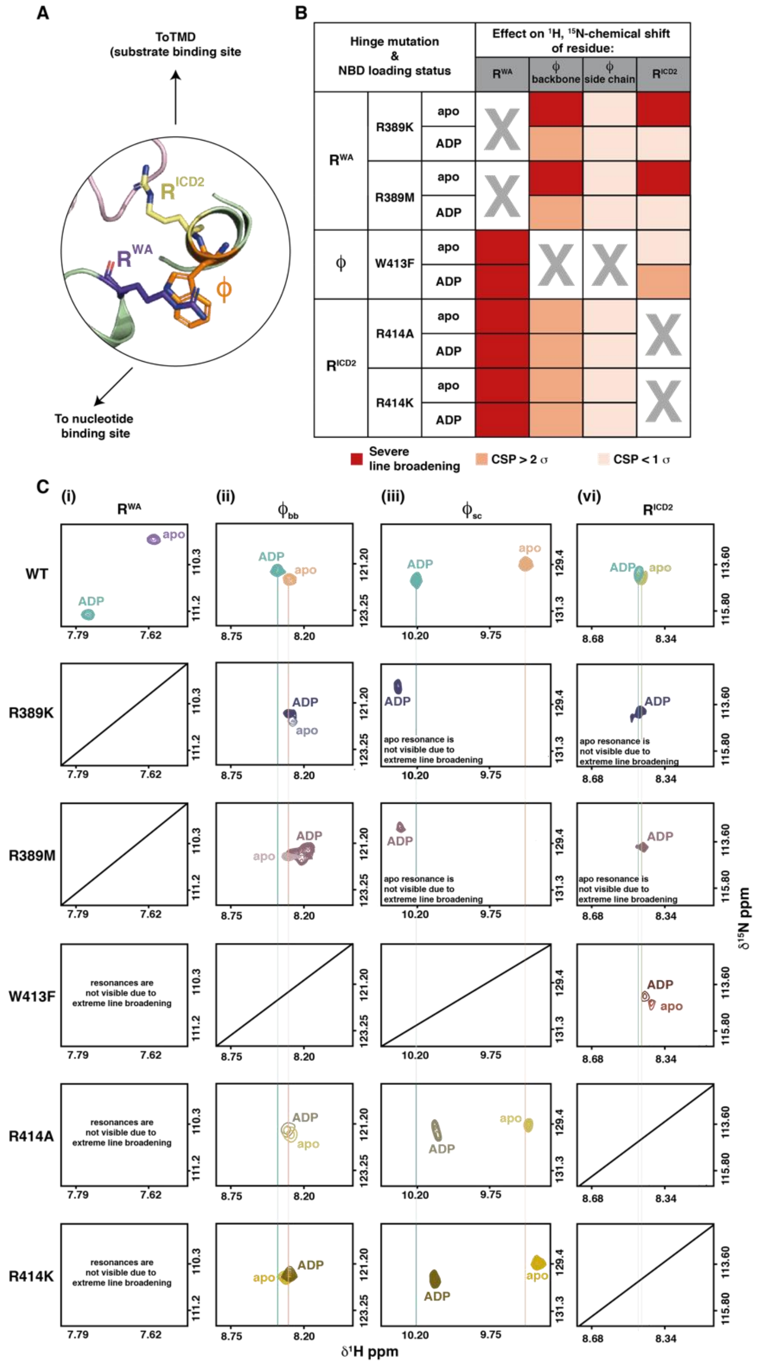


**Supplementary Figure 4: The hinge residues are structurally and dynamically coupled.** (**A**) Cartoon of communication hinge highlighting relative position of the three conserved residues in BmrA. (**B**) Table summarizing the data shown in (C), i.e. the nucleotide dependent effects on NMR chemical shift and line width for the $^{1}$H, $^{15}$N resonances of the hinge residues in the $^{15}$N-labeled BmrA NBD, upon mutating one of the three hinge residues. The respective mutated residue is crossed out. (**C**) Zoom into the $^{1}$H, $^{15}$N-HSQC to show the NH resonances for the three hinge residues in the WT NBD (top row) and upon mutating each of the hinge residues individually (bottom rows). Chemical shifts of WT NMR in the apo state and in the presence of 10 mM ADP are indicated shown as colored lines. Note that for residue $\phi$, a tryptophan, both the backbone and sidechain indole NH resonances can be observed (denoted $\phi_{bb}$ and $\phi_{sc}$, respectively). The consequences of mutating either of the hinge residues on the chemical shift and linewidth compared to the WT is summarized in the table shown in (B).

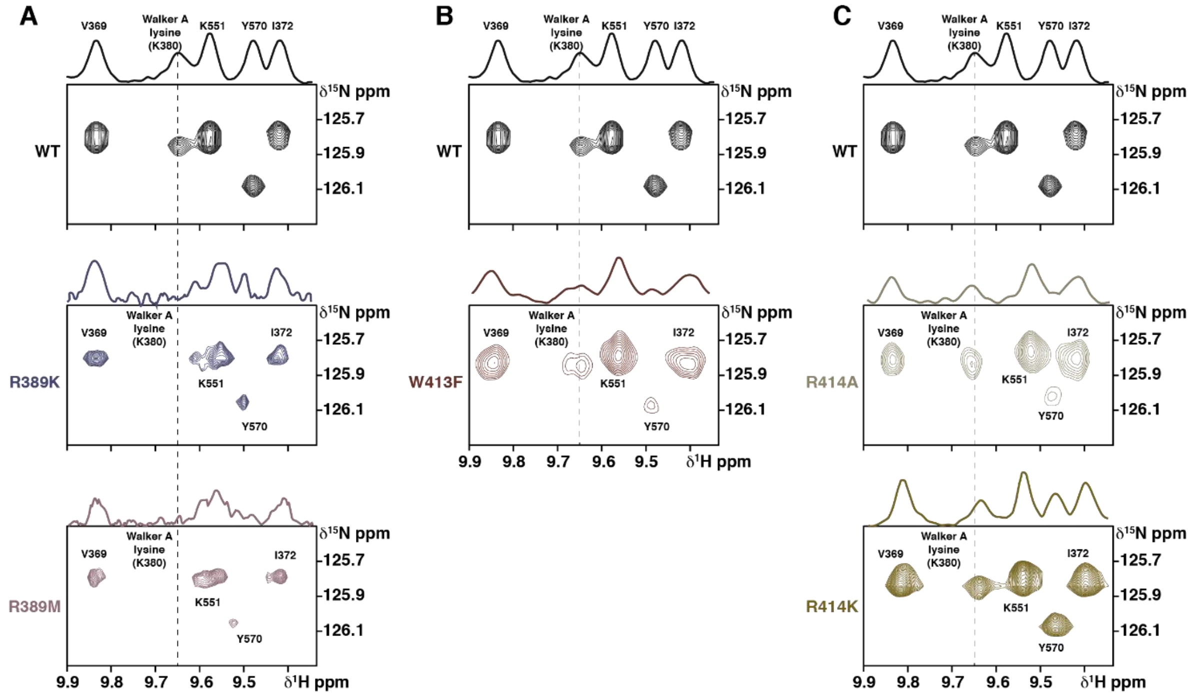


**Supplementary Figure 5: Effects of hinge mutations on the conserved Walker A lysine in the nucleotide binding site.** (**A, B, C**) Regions from the $^{1}H$, $^{15}N$ HSQC NMR spectra of BmrA NBD WT and hinge mutants, focusing on the resonance of the conserved Walker A lysine K380, which becomes visible only in the presence of ADP. Thus, all spectra were recorded with 10 mM ADP at 298K on a 600 MHz spectrometer equipped with a cryogenic triple resonance probe (Bruker GmbH, Karlsruhe, Germany, using a 200 µM sample. For better visibility, the 1D projections are shown on top of each spectrum, and the $^{1}H$ chemical shift of the K380 resonance in the WT NBD is marked with a dashed line. Upon mutation of residue $R^{WA}$ (A), $\phi$ (B) or $R^{ICD2}$ (C), chemical shift changes and line broadening for K380 occurred, showing that nucleotide binding site and hinge are structurally and dynamically coupled.

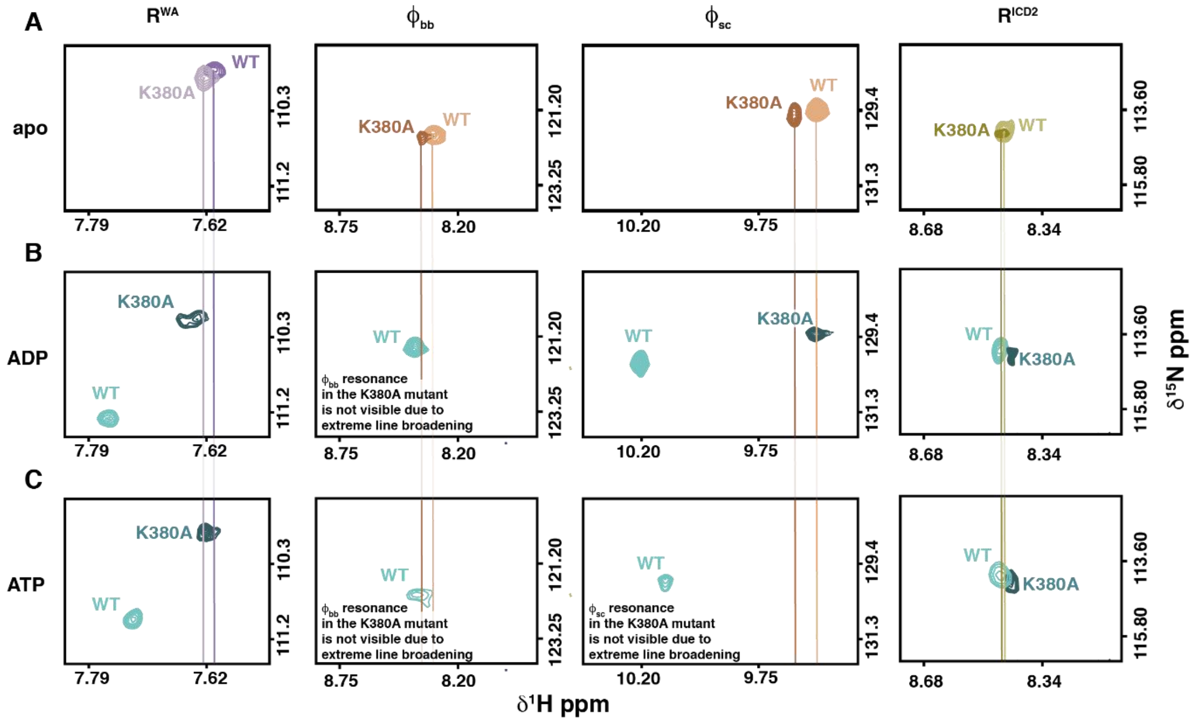


**Supplementary Figure 6: Effect of Walker A mutant K380A on hinge residues.** (**A**) Comparison of chemical shifts of hinge residues between BmrA NBD WT and Walker A K380A mutant in the absence of nucleotides showing that mutation of the nucleotide binding site alone is sensed by the hinge residues. (**B, C**) Comparison of chemical shifts of hinge residues between BmrA NBD WT and Walker A K380A mutant in the presence of 10 mM ADP (B) and ATP (C). In some cases, mutation of the Walker A residue led to severe line broadening in the hinge. All spectra were recorded on a 600 MHz spectrometer equipped with a cryogenic triple resonance probe (Bruker GmbH, Karlsruhe, Germany) at 298K using a 200 µM sample.

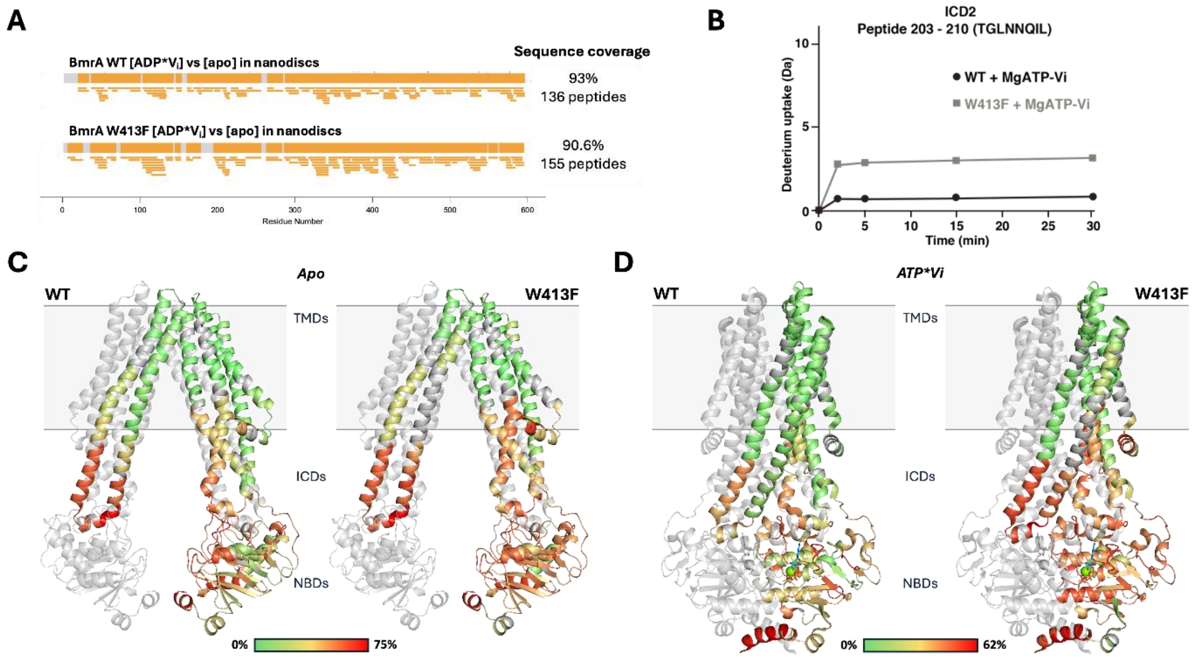


**Supplementary Figure 7. Impact of BmrA hinge W413F mutation on transporter dynamics in lipid nanodiscs followed by Hydrogen Deuterium Exchange coupled with mass spectrometry.** (**A**) Sequence coverage map of BmrA WT (upper panel) and W413F (lower panel) showing peptides (orange bars) shared between the apo- and the MgADP*Vi trapped states (upon incubation with 10 mM $Mg^{2+}$, 10 mM ATP and 1 mM Vi). (**B**) Comparison of deuterium uptake for a peptide (residues 203–210 (TGLNNQIL)) within ICD2 between BmrA WT (black) and the hinge mutant (gray) after incubation with MgATP-Vi exhibits reduced HDX in the MgADP*Vi trapped state for the WT, relative to the hinge mutant. (**C**) HDX after 30 min deuteration for BmrA WT (left) and hinge mutant (right) in the apo state. HDX was plotted on the cryoEM structure of BmrA in the open inward state (PDB: 8QOE). For clarity, one BmrA monomer was kept transparent, while the second monomer was colored in a deuteration scale ranging from 0 to 75%, with 75% representing the maximum deuteration observed under these conditions. Uncovered peptides are represented in dark gray. (**D**) HDX after 30 min deuteration for BmrA WT (left) and hinge mutant (right) in the MgATP*Vi trapped state plotted on the cryoEM structure of BmrA in the OF state (PDB: 7OW8). One BmrA monomer is transparent, while the second monomer is colored in a deuteration scale ranging from 0 to 62%, with 62% representing the maximum deuteration observed under these conditions.

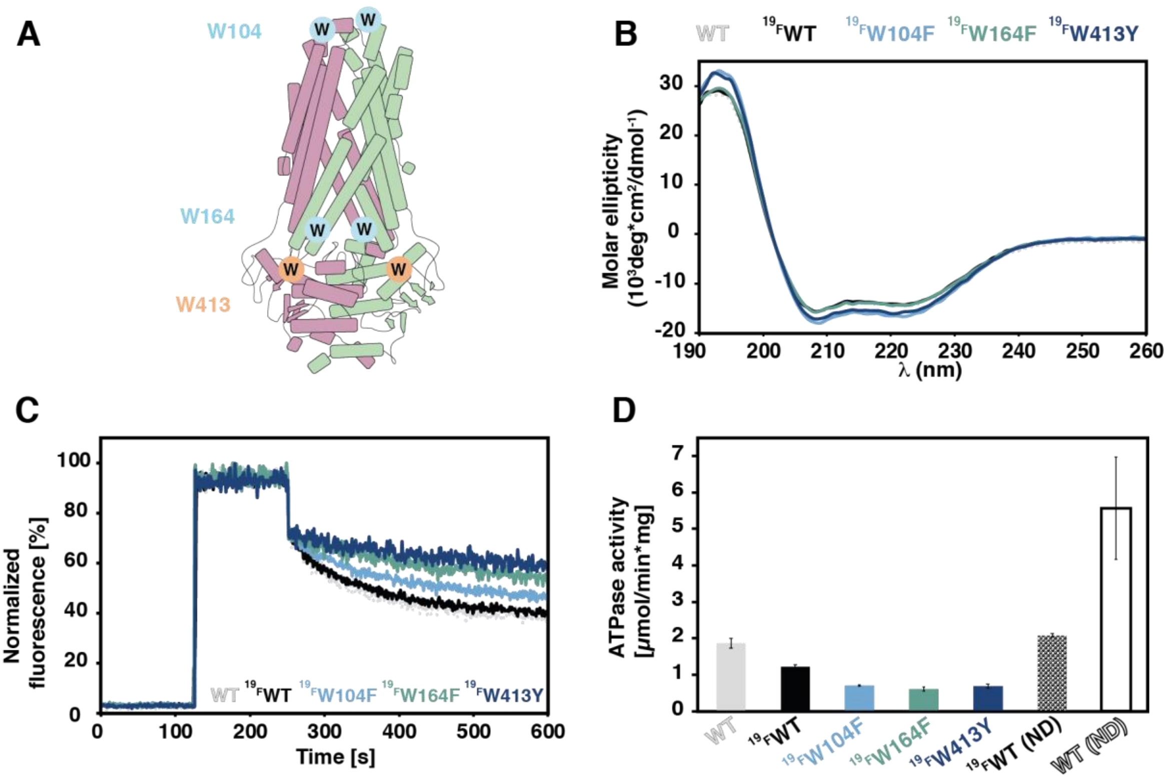


**Supplementary Figure 8: Fluorination of tryptophan residues in full-length BmrA does not have a major impact on structural and functional integrity.** (**A**) Position of the three native tryptophan residues in BmrA. Note that W413 is the only tryptophan residue in the NBD and part of the hinge (residue ϕ, orange).(**B**) Circular dichroism spectra of unlabeled BmrA WT (light grey), $^{19}$F-Trp labeled WT (black) and the three labeled single point mutants created to assign the $^{19}$F NMR spectra shown in Fig. 3E. (**C**) Fluorescence-based transport assay with doxorubicin in inside out vesicles prepared from cells overexpressing BmrA variants. All traces normalized to the unlabeled WT curve (light grey). (**D**) ATPase activity of BmrA variants in DDM/NaCholate (first five bars) or reconstituted in MSP1E3D1 nanodiscs prepared with *E. coli* polar lipid extract (last two bars on right, signified by 'ND'). Importantly, fluorinated WT BmrA reconstituted in nanodiscs retains full activity compared to the unlabeled WT in detergent micelles.

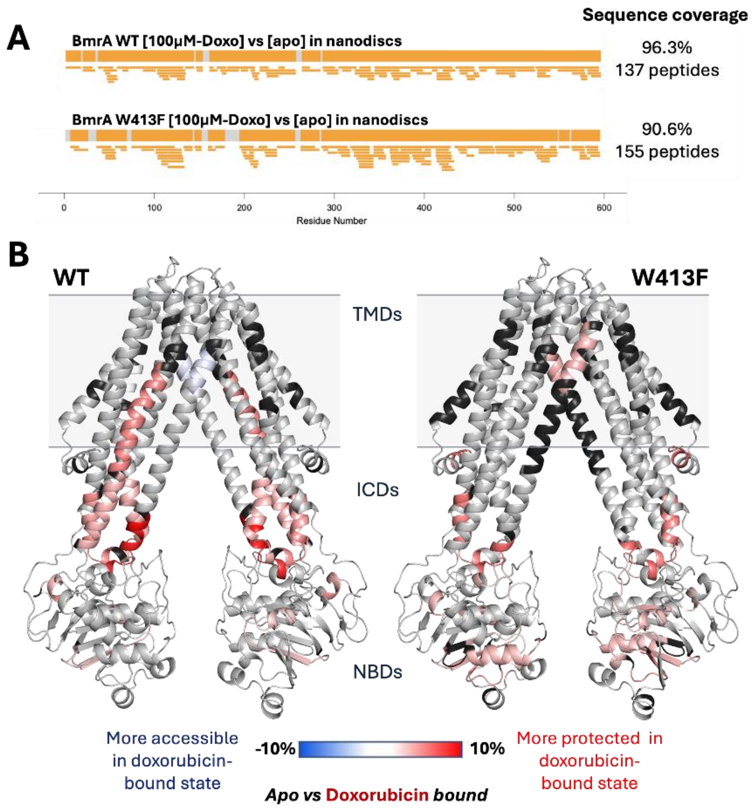


**Supplementary Figure 9: Hydrogen Deuterium Exchange coupled with mass spectrometry on BmrA in lipid nanodiscs in the presence of doxorubicin.** (**A**) Sequence coverage map of BmrA WT (upper panel) and W413F (lower panel) showing peptides (orange bars) shared between the apo- and the doxorubicin-bound states (upon addition of 100 µM doxorubicin and incubation for a minimum of 15 min at 20°C). Similar peptide coverage as for the apo and ATP*$V_i$ trapped transporter was achieved. (**B**) HDX-MS in the absence and presence of doxorubicin. Shown are the differences between the apo and doxorubicin-bound states for BmrA WT (left) after 15 min deuteration, plotted on the cryoEM structure of BmrA in the inward-facing state (PDB: 8QOE). The same experiment was repeated with the hinge mutant W413F (right), which resulted in noticeably reduced effects on the HDX in the presence of the drug. Peptides with no significant deuteration differences (p-value = 0.05, see Methods) are shown in light gray and uncovered peptides in black.

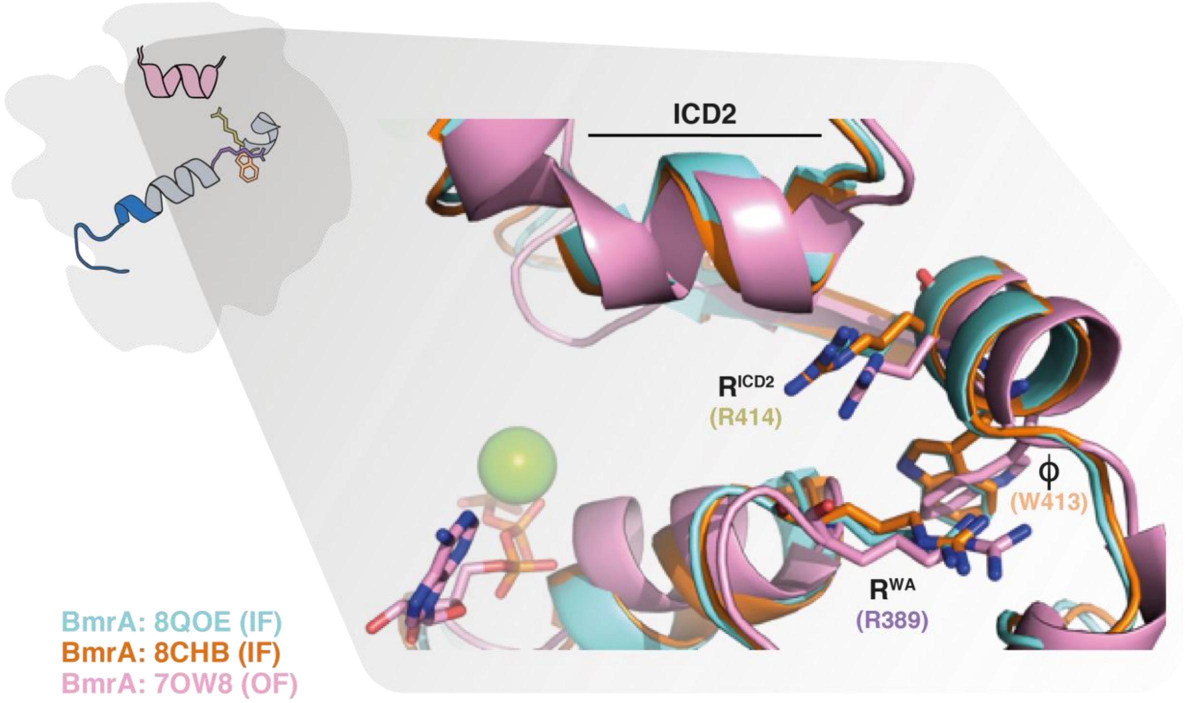


**Supplementary Figure 10: Comparison of the hinge region in the inward and outward facing conformations of BmrA.** Overlay of the hinge region in three available structures of BmrA. Two structures were solved in the inward facing (IF) conformation (Di Cesare et al., 2024) (PDBs: 8QOE, 8CHB) and the third structure was determined in the outward facing (OF) conformation bound to MgATP using the catalytically inactive mutant E504A (Chaptal et al., 2022) (7OW8). These three structures were selected after visual inspection of the electron densities in the six existing structures of BmrA available in the PDB at this site, allowing the proper positioning of the side chains of the hinge residues in the modelled structures. In the three structures, the $R^{WA}$, $\phi$ and $R^{ICD2}$ side chains occupy highly similar positions in agreement with the observed electron densities, resulting in a highly similar side chain orientation of residue $R^{ICD2}$ pointing towards the coupling helix of the protomer in *trans*.
